# Principles of NMDA receptor co-agonism at cortical fast-spiking GABAergic interneurons in the adolescent prefrontal cortex

**DOI:** 10.64898/2026.08.30.747826

**Authors:** Isis N. O. Souza, Pierre Lecouflet, Zeynep Okur, Steeve Maldera, Brigitte Potier, Loredano Pollegioni, Peter Scheiffele, Jean-Pierre Mothet

**Author notes:** Corresponding author: Jean-Pierre Mothet. **Email:**. These authors contributed equally to this work (Isis N. O. Souza, Pierre Lecouflet).

## Abstract

N-methyl-D-aspartate receptors (NMDARs) populate fast-spiking (FS)-parvalbumin-positive (PV^+^) GABAergic interneurons (INs), where they play a critical role in shaping circuit motifs and memory. However, it is largely unknown whether and how NMDARs at FS-PV^+^-INs are gated by their co-agonists and the functional relevance of such modulations for their synaptic coupling with excitatory neurons. Here, we report that FS-PV^+^-INs in the adolescent mouse prefrontal cortex, an area central to complex cognitive operation exhibit functional GluN2B/D containing NMDARs. These receptors contribute to the excitatory drive of FS-PV^+^-INs and to the feedforward inhibition, controlling short-term and long-term synaptic plasticity. While the identity of the co-agonist controlling GABAergic tone is tuned by the synaptic activity regime from d-serine to glycine, we reveal that it remains largely unchanged at the excitatory synapse with d-serine being the sole co-agonist gating NMDARs. Lastly, we show that d-serine-deficient mice, a model of NMDAR hypofunction show selective attenuation of PV^+^-INs excitation together with selective loss of temporal summation and long-term plasticity at the excitatory synapse. Our study reveals the segregation of pools of NMDARs at the soma and dendrites that are differently sensitive to D-serine or glycine, the existence of distinct modes of activity-dependent regulation of these NMDARs by their co-agonists at this major type of GABAergic INs, and hence the rules governing cortical inhibition by FS-PV^+^-INs during a critical period of late postnatal development.

**Significance Statement:** Our study sheds new light on brain circuit’s physiology by uncovering the general principles of NMDARs regulation at GABAergic interneurons by upstream signals released in their surrounding microenvironment. We discovered that d-serine but not glycine is critical for maintaining the activity of NMDARs at FS-PV^+^ interneurons in the prelimbic area of the prefrontal cortex and that loss of its functions would recapitulate the synaptic deficits observed in several neuropsychiatric diseases like schizophrenia. These results could be exploited for the future development of more effective clinical interventions for treating NMDAR hypofunction by targeting inhibitory neurons.

## Introduction

GABAergic interneurons are in charge of the dynamic adjustment of the level of excitation insured by glutamatergic neurons to maintain a proper excitation/inhibition balance, and are therefore essential for information processing within the brain (1, 2). The fast-spiking parvalbumin-containing interneurons (FS-PV^+^ INs) represent the largest population of cortical inhibitory neurons (1) and, by providing the major perisomatic inhibitory drive to excitatory pyramidal neurons (PNs), play a critical role in circuit physiology (3, 4). As a consequence, malfunction of these GABAergic interneurons is central to several neuropsychiatric disorders (5–7).

FS-PV^+^-INs are primarily recruited by glutamatergic synaptic input from the PNs. In particular, FS-PV^+^-INs express N-methyl-D-aspartate receptors (NMDARs) (3, 4) that are pivotal for circuit entrainment and for generating network oscillations that encode numerous cognitive processes. In particular, NMDARs play a critical role in the integration and maturation of GABAergic interneurons in functional circuits during critical periods? of postnatal development. Selective ablation of NMDARs during early postnatal development results in synaptic dysconnectivity with PNs and in corrupted network oscillations and behavioural deficits (8–10). Thus, hypofunction of NMDAR onto FS-PV^+^-INs has emerged as a cause of various neurodevelopmental disorders including schizophrenia (SCZ) and autism spectrum disorders (ASD) (10, 11). However, how NMDAR at FS-PV^+^-INs are physiologically regulated by extracellular signals is unknown. NMDARs are heterotetrameric molecular structures typically composed of two GluN1 subunits and two GluN2 subunits (12, 13). Yet, activation of the canonical NMDARs requires simultaneous binding of _L_-glutamate and a co-agonist, glycine or d-serine (12–14). Paradoxically, more than two decades after the discovery of the action of glycine and d-serine at NMDAR (15–17), we are still ignoring whether and how the activity of specific populations of GABAergic interneurons and their synaptic connections with PNs are regulated by NMDAR co-agonists, and the physiopathological relevance of such modulation particularly during critical periods of development remains unaddressed. Furthermore, d-serine deficient mice were previously shown to recapitulate a spectrum of symptoms relevant to SCZ (18–20) with suspicious FS-PV^+^ IN dysfunction (21) but the underlying mechanisms remain poorly understood. In this study, we investigated the physiological conditions that specify d-serine and glycine co-agonism of NMDARs at FS-PV^+^-INs during adolescence, focusing on the prelimbic area (PrL) of the prefrontal cortex (PFC), a late-maturing region central to complex cognitive operations (22–24) and one where FS-PV^+^-IN dysfunction has invariably been implicated in neurodevelopmental disorders including SCZ (25). We show that adolescent FS-PV^+^-INs retain functional somatodendritic NMDARs containing GluN2B and GluN2C/D subunits, which shape both the excitatory drive they receive and the GABAergic output they generate. Rather than a uniform dependence on a single co-agonist, we find that co-agonist identity is governed by synapse-specific and activity-dependent rules: at the excitatory PN-*to*-FS-PV^+^-IN synapse, d-serine remains the dominant, largely non-substitutable co-agonist that gates NMDARs across the activity regimes we tested, whereas at the inhibitory FS-PV^+^-IN-*to*-PN synapse, the co-agonist requirement shifts with synaptic activity, with glycine becoming increasingly able to sustain NMDAR function as activity intensifies. This functional segregation raises the possibility that distinct pools of NMDARs, differentially distributed across the somatic and dendritic compartments of FS-PV^+^-INs, show differential sensitivity to d-serine *versus* glycine. Consistent with this asymmetry, FS-PV^+^-INs from d-serine-deficient mice show attenuated firing and a selective loss of excitatory long-term potentiation (eLTP), while temporal summation and inhibitory long-term potentiation (iLTP) at their output synapses onto PNs are preserved — indicating that glycine can act as a partial, pathway-specific substitute for d-serine rather than a general compensatory mechanism. Together, these findings refine the current view of NMDAR hypofunction at GABAergic interneurons in psychiatric disease, revealing distinct, compartment- and activity-dependent modes of co-agonist regulation that differentially control the excitatory input and the feedforward inhibitory output of FS-PV^+^-INs. They also broaden the physiological landscape of d-serine signaling in the brain and point to more selective, pathway-informed strategies for therapeutic intervention.

## Results

### Functional GluN1-GluN2 NMDARs are present at FS-PV^+^-INs at late adolescence

We focused on the regulation of FS-PV^+^ INs from layer 5 of the prelimbic area (PrL) of the prefrontal cortex during late adolescence, a critical period for the onset of clinical symptoms for many psychiatric disorders in humans (26). These cells can be readily identified in PV^+^-*td*Tomato fluorescent reporter P45-60 mice (Fig. 1a). Upon injection of depolarizing current pulses, these interneurons exhibited typical high frequency firing patterns and non-adaptive spikes with short lasting action potentials (APs) (half-width 0.51 ± 0.06 ms) and large fast after-hyperpolarization (AHPs, 21.48 ± 4.03 mV) (Supplementary Fig. 1). Although functional NMDARs are known to populate different classes of GABAergic interneurons during early cortical development (27, 28), their presence in adolescent and adult FS-PV^+^ INs and their potential to modulate FS-PV^+^ INs bursting activity remain highly debated (29–32). Bath applied NMDA (10 µM) consistently increased the firing activity of FS-PV^+^ INs in response to depolarizing current pulses (Supplementary Figs. 2&3), an effect which is systematically nulled by including the non-competitive NMDAR antagonist MK801 (2-3 mM) in the intracellular solution of the recording pipette (iMK801) (30, 31). More importantly, iMK801 in the absence of NMDA consistently reduced the excitability of the FS-PV^+^ INs (Fig. 1b), suggesting that at least a subset of NMDARs is tonically activated. Indeed, bath application of the selective NMDAR antagonist d-AP5 (25 µM) induces a negative shift in the holding current as well as synaptic noise reduction (Fig. 1c). We next addressed for the presence of synaptic NMDARs by recording electrically evoked EPSCs. NMDA EPSCs were pharmacologically isolated using blockers of AMPA receptors (NBQX, 20μM), GABA_A_ receptors (picrotoxin, 50μM), and strychnine sensitive glycine receptors (strychnine, 1 µM) in normal Mg^2+^ (1.5 mM) ACSF. Electrical stimulation (0.1 Hz) elicited typical current-voltage (I-V) curve of NMDA EPSCs (Fig. 1d) that reversed at 0 mV and showed inward rectification. iMK801 selectively blocked NMDA EPSCs while preserving AMPA EPSCs (Fig. 1d). Noteworthy, the NMDAR/AMPAR peak amplitude ratio was ∼1.0, indicating that FS-PV^+^ INs in late adolescent mice still express large fraction of functional NMDARs as reported recently (32, 33). High-resolutive immunogold labelling of freeze-fracture replica reveals that GluN1 subunits which reflects the presence of NMDARs are substantially expressed in the somadendritic arbor of the FS-PV^+^ INs while GluN1 is absent from axonal boutons (Prof Z Nusser, personal communication). Therefore, and in contrast with most studies (32–34), operational NMDARs largely populate FS-PV^+^ INs beyond early postnatal development, where they significantly contribute to modulate the firing and synaptic activities of that class of inhibitory interneurons. The GluN2C/2D subunits are preferentially expressed in GABAergic interneurons in the mature brain (28, 35, 36). Using Fluorescent *in situ* hydridization (FISH), we confirmed that GluN2D is enriched in FS-PV^+^ INs at P50-55, showing a similar number of puncta to GluN2B (Fig. 1e, Supplementary Fig 4). Accordingly, bath application of NAB-14 (20 µM), a potent and selective negative allosteric modulator of GluN2C/2D-containing NMDARs, significantly reduced NMDA EPSCs peak amplitude (Fig. 1f). Likewise, bath applied Ro-25-6981 (4 µM), a selective blocker of GluN2B-containing NMDARs further reduced the NMDA EPSCs in a similar fashion (Fig. 1f). Taken together, these data indicate that post-synaptic putative GluN1/2B/2D NMDARs are present at FS-PV^+^ INs (33) during adolescence and function to control synaptic activity and intrinsic excitability of these interneurons.

**Figure 1.**
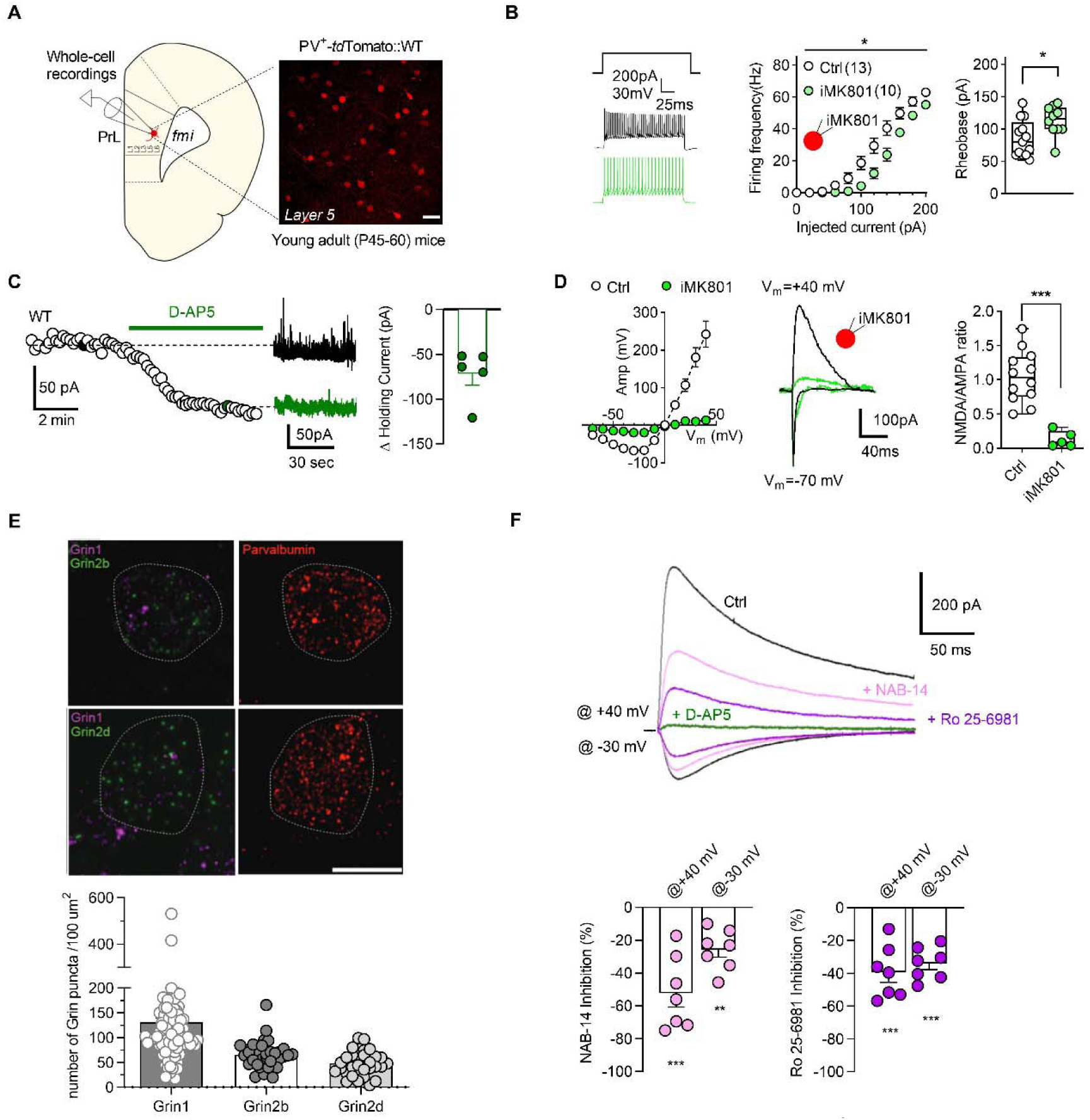
Functional GluN2 containing NMDARs populate FS-PV^+^ interneurons at late adolescence. (**A**) Left, schematic diagram depicting a mPFC slice from which layer 5 FS-PV^+^ INs were recorded in PV^+^-*td*Tomato::WT mice. Right, representative confocal image of *td*Tomato expressing interneurons in layer 5 of the prelimbic area (PrL) of P45-70 mouse. Scale bar: 40 μm. (B) Action potentials recordings (Left) before (black trace) and after inclusion of MK801 in the patch pipette (iMK801, light green) in FS-PV^+^ INs. Spike frequency (middle) from graded current injections (0 to 200 pA in 20-pA steps) on interneurons with or without iMK801 (n= 10 cells, *p*=0.0118, F(1,21)= 7.603, two-way ANOVA) compared to Ctrl (n= 13 cells). Right, boxplot graphs showing the increase of rheobase with iMK801 compared to Ctrl (*p*= 0.018, Mann-Whitney test). (C) Tonic current recorded at +40 mV (Left) in FS-PV^+^ INs before (black trace) and after bath application of the NMDAR blocker, D-AP5 (green trace). Average holding current reduction analyzed by One sample Wilcoxon test (*p*=0.0625). (**D**) Current-voltage (I-V) relationship showing the inhibition of NMDA EPSCs amplitude by iMK801 in FS-PV^+^ INs (n= 5 cells, *p*=0.0045, F(1,15)= 11.13, two-way ANOVA) compared to control (n= 12 cells). Examples of NMDA EPSCs (+40 mV) and AMPA-EPSCs (−70 mV) obtained in Ctrl condition (black trace) and in the presence of iMK801 (light green trace). Note that AMPA EPSC are not affected by iMK801. Box plots (right) show NMDA/AMPA ratio (n= 5 cells, *p*=0.0003, Mann-Whitney test). (**E**) Fluorescent *in-situ* hybridization (FISH) micrographs indicated the presence of Grin1, Grin2b and Grin2d RNA in FS-PV^+^ INs from WT mice. Scale bar: 10 µm. Bar graphs represent the number of Grin puncta/100 µm^2^. (**F**) Left, representative NMDA EPSCs traces recorded in control conditions (black), NAB-14 (mauve), Ro 25-6981 (violet) or D-AP5 (green). Right, histograms show that Ro 25-6981 (n= 7 cells) and NAB-14 (n= 7 cells) inhibit similarly NMDA EPSCs at FS-PV^+^ INs from WT mice when recorded at either −30 or +40mV. NAB-14 @ +40 mV, *p*=0.0156; NAB-14 @-30 mV, *p*=0.0156; Ro 25-6981 @ +40 mV, *p*=0.0156; Ro 25-6981 @ −30 mV, *p*= 0.0156 (One sample Wilcoxon test). \**p*< 0.05, \*\**p*< 0.01, \*\*\**p*< 0.001. Data are expressed as mean ± SEM.

### Gating of NMDARs by d-serine selectively controls the intrinsic excitability and the excitatory drive of FS-PV^+^-INs

Conventional GluN1-GluN2 containing NMDARs require a co-agonist, d-serine or glycine for their activation (12–14). We next wondered whether glycine and/or d-serine may regulate NMDARs at the FS-PV^+^ INs. Boosting extracellular glycine levels by blocking the glycine transporter GlyT1 with ALX5407 (1 µM) (37–40) did not alter spike frequency of FS-PV^+^ INs (Fig. 2b, Supplementary Fig. 3). Similarly, depleting extracellular glycine with the enzymatic scavenger *Bs*GO (0.2 U/mL) (38–40) did not alter the firing activity of FS-PV^+^ INs (Fig. 2c, Supplementary Fig. 3). An effect of *Bs*GO reflecting a decrease in glycine levels became evident at injected currents >200 pA (data not shown). These findings suggest that ambient extracellular glycine is normally low and does not regulate firing activity of FS-PV^+^ INs. Conversely, favoring the presence of endogenous d-serine by bath-application of CBIO (1 µM) (40–42), an inhibitor of the d-serine degrading enzyme d-amino acid oxidase (DAAO) (14, 43–46), increased the excitability of FS-PV^+^ INs (Fig. 2d, Supplementary Fig. 3), but remarkedly not of PNs (Supplementary Fig. 5). iMK801 impeded the potentiating action of CBIO (Fig. 2e, Supplementary Fig. 3) thus demonstrating that the modulation of firing activity does not result from network activity and is most likely caused by a direct action of d-serine on NMDARs at the FS-PV^+^ INs. Altogether, these findings support the view that the intrinsic activity of FS-PV^+^ INs highly depends on the strict occupancy of NMDAR co-agonist binding site by d-serine and that these NMDAR are tonically activated under basal conditions. We then wondered whether the excitatory synaptic PN-*to*-PV^+^ coupling might also be determined by d-serine. As before, NMDA EPSCs were pharmacologically isolated using NBQX (20μM), picrotoxin (50μM) and strychnine (1 µM). Selective and acute depletion of d-serine with the enzymatic scavenger *Rg*DAAO (0.2 U/mL) (38–40, 47) did not affect the voltage-dependency of the NMDA EPSCs (Fig. 2f) but yielded to a ∼40% reduction in the NMDAR/AMPAR ratio tied to its specific action on NMDA EPSCs (Fig. 2g). Noteworthy, *Rg*DAAO also affected the kinetics of NMDA EPSCs by reducing the rising time without impacting the decay time suggesting that modalities of NMDARs gating have been changed upon removal of d-serine (Supplementary Fig 6). As shown in Fig. 2g, NMDA EPSCs persisted in the presence of *Rg*DAAO, suggesting that glycine might also contribute to regulate synaptic NMDARs (38–40). However, application of *Bs*GO (0.2 U/ml) (38–40) to acutely remove glycine neither affected NMDA EPSCs peak amplitude nor current-voltage curve (Fig. 2f,g). Surprisingly, *Bs*GO failed to abolish the remaining d-AP5 sensitive NMDA EPSCs observed under *Rg*DAAO (Fig. 2g, condition x2), further demonstrating that glycine under normal conditions does not gate synaptic NMDARs at FS-PV^+^ INs and that the partial inhibitory action of *Rg*DAAO reflects an incomplete ablation of the extracellular d-serine as already reported. As expected, CBIO, which elevates synaptic levels of d-serine, increased NMDA EPSCs (Fig. 2h), suggesting that the co-agonist site of synaptic NMDAR on FS-PV^+^ INs was not saturated. Intriguingly, blockade of GlyT1 with ALX5407, which increases ambiant glycine levels, increased the amplitude of NMDA-EPSCs as well (Fig. 2h). CV^-2^ analysis revealed that CBIO and ALX5407 exerted their potentiating effects purely at postsynaptic NMDARs (Supplementary Fig. 7). In conclusion, these results revealed that NMDAR at FS-PV^+^ INs in late adolescence are primarily gated by d-serine and not glycine at low synaptic activity regime and point that glycine transporters (e.g GlyT1) are efficiently maintaining the levels of glycine below effective concentrations within the synaptic cleft, as already observed at excitatory synapses between PNs in the hippocampus (38, 40) or amygdala (39).

**Figure 2.**
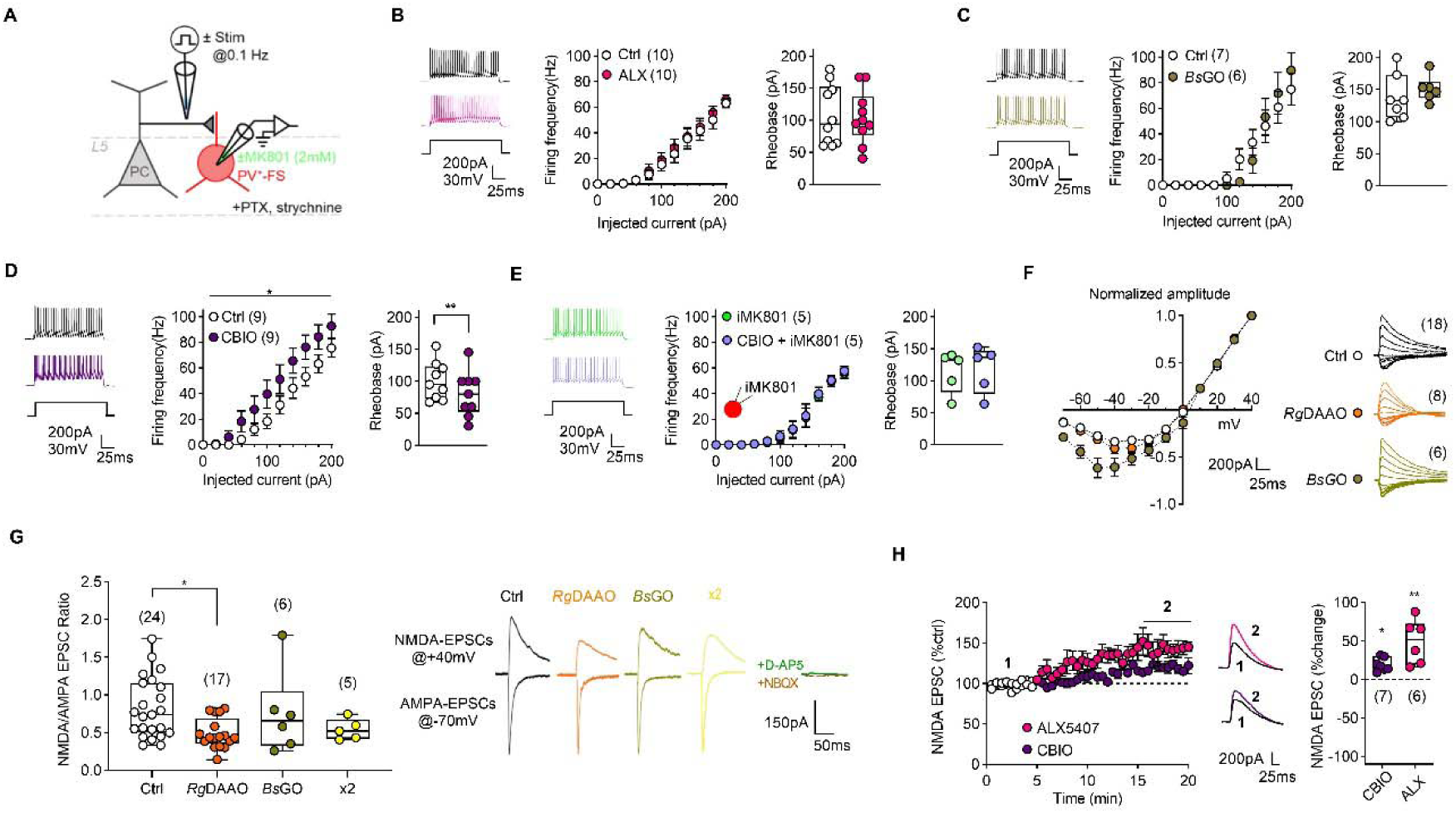
NMDAR regulation of intrinsic excitability and strength of excitatory input depends unrequitedly on d-serine co-agonism. (**A**) Schematic diagram showing the placement of recording electrode in mPFC layer 5, where FS-PV^+^ INs were recorded in PV^+^-*td*Tomato::WT mice, with bath application of PTX and strychnine (all panels). For isolating NMDA EPSCs NBQX was added (panels F to H). (**B** and **C**) Active potential recordings (Left) of *td*Tomato-positive FS-PV^+^ interneurons in the mPFC in the absence (Ctrl) or in the presence of ALX-5407 (1 µM, fuschia) or when slices were treated with *Bs*GO (0.4 U/ml, olive) to manipulate glycine levels. Spike counts (middle) from graded current injections (0 to 200 pA in 20-pA steps) on interneurons with varying conditions. ALX-5407: n= 10 cells, *p*= 0.310, F(1,9)= 1.158, two-way ANOVA. *Bs*GO: n= 7 control and 6 *Bs*GO-treated cells, *p*= 0.961, F(1,11)= 0.002, two-way ANOVA. Box plots (right) show the effects of different treatments on rheobase. Data were analyzed with Wilcoxon test (ALX-5407, *p*= 0.434; *Bs*GO, *p*= 0,281). (**D** and **E**) Active potential recordings (Left) of *td*Tomato-positive FS-PV^+^ interneurons in the mPFC in the absence (Ctrl) or in the presence of CBIO (1 µM, purple) alone or when MK801 was introduced (light green) in the FS-PV^+^ interneuron. Spike counts (middle) from graded current injections (0 to 200 pA in 25-pA steps) on interneurons with varying conditions. CBIO: n= 9 cells, p= 0.012, F(1,8)= 10.340, two-way ANOVA). CBIO+iMK801: n= 5 cells, p= 0.209, F(1,4)= 2.233, two-way ANOVA). Box plots (right) show the effects of the different treatment on rheobase. Data were analyzed with Wilcoxon test (CBIO *vs* Ctrl, *p*=0.004; CBIO *vs* CBIO+ iMK801, *p*> 0.999). (**F**) I-V relationship of normalized peak amplitude of NMDA EPSCs (Left) recorded from −70 to +40 mV (in 10 mV steps) in WT mice (white), in the presence of the enzymatic scavenger *Rg*DAAO (orange, n= 12 Ctrl and 7 *Rg*DAAO-treated cells, *p*= 0.134, F(1,17)= 2.474, two-way ANOVA) or with *Bs*GO (n= 6 control and 6 *Bs*GO-treated cells, *p*= 0.121, F(1,10)= 2.872, two-way ANOVA). Right, sample traces of NMDA EPSC under different conditions and for each voltage potential. (**G**) Summary box plot (left) of the NMDA/AMPA peak amplitude ratio in control (Ctrl, n=24 cells), or when slices were treated either with *Rg*DAAO (Orange, 17 cells), *Bs*GO (Olive, n=6 cells) or both (x2, yellow, n=5 cells). NMDA EPSCs were recorded at +40 mV and AMPA EPSCs at −70 mV. Ctrl *vs Rg*DAAO *p*= 0.0273, *Rg*DAAO *vs Bs*GO *p>* 0.999, *Bs*GO *vs* x2 *p>* 0.999, Kruskal-Wallis test with post-hoc Dunn’s test. Right, sample traces of EPSCs. Note that NMDA EPSCs and AMPA EPSCs were nulled by bathing D-AP5 (dark green, n=4 cells) or NBQX (Brown, n=4 cells), respectively. (**H**) Time course (left) showing the averaged effect of bath applied CBIO and ALX-5407 on NMDA EPSCs. Histogram (right) shows the relative changes induced by both compounds on the NMDA EPSC peak amplitude (CBIO: 7 cells, *p*= 0.016; ALX5407 n= 6 cells, *p*= 0.031, Wilcoxon test). \**p*< 0.05, \*\**p*< 0.01. All data are expressed as mean ± SEM.

### Acute loss-of-function of d-serine impairs excitatory temporal summation and synaptic plasticity onto FS-PV^+^ Ins

High-frequency dependent synaptic and network activities in the mPFC correlates with cognitive performance. We next explored whether the identity of the co-agonist driving NMDAR at FS-PV^+^ INs might be determined by the level of synaptic activity regime (39, 40). To do so, we first analyzed the role of d-serine in controlling NMDAR-dependent temporal summation, a form of activity-dependent short-term plasticity (Fig. 3) with realistic trains of stimulation. The 2, 20 or 50 Hz trains are typical of mPFC FS-PV^+^ INs firing rates during attentional processing (48–50). At 2 and 20Hz (Fig. 3a), individual NMDA EPSCs were clearly discernible for each pulse of stimulus current and exhibited a clear amplitude attenuation (synaptic fatigue) which was precipitated by increasing frequencies (51, 52). Acute d-serine depletion by treating the slices with *Rg*DAAO accelerated and increased the depression rate at 2 Hz but not at 20 Hz nor at 50Hz in comparison to control slices. At 50Hz, NMDAR EPSCs were not discernible and exhibited temporal summation of the first three trains before substantial depression appeared (Fig. 3a). Such pattern is caused by the slow kinetics of the NMDAR channels (12, 13) thus occluding single NMDA EPSCs to return to baseline before the next stimulation. The presence of a threshold level of d-serine is necessary for NMDAR to maintain persistent states of activity at θ frequencies (∼2Hz). These data show that the synaptic depression became independent of the levels of the co-agonist d-serine at β and γ ranges where glycine most likely can be recruited. Finally, although the excitatory PN-*to*-PV^+^ synapse has been shown to undergo various forms of activity-dependent long-term plasticity (53–58), whether NMDAR-dependent long-term potentiation (LTP) of such synapses relies on a specific co-agonist is still unknown. High-frequency stimulation (HFS) with postsynaptic depolarization of the FS-PV^+^ IN to 0 mV invariably induced a d-AP5-sensitive excitatory LTP (eLTP, Fig. 3b). This potentiation appeared not to depend on somatodendritic voltage-gated Na^+^ channels activation, since the blocker QX314 (5 mM) was included in the patch pipette. The mean paired pulse ratio (PPR) of evoked EPSCs (interstimulus interval 30 ms) was not significantly changed upon this Hebbian eLTP at FS-PV^+^ INs (Supplementary Fig. 8a). In addition, analysis of CV^-2^ confirmed the postsynaptic locus of the eLTP (Supplementary Fig. 8b). These results suggest that this form of eLTP is expressed with no apparent change in presynaptic function (e.g transmitter release). More importantly, the same protocol failed to induce eLTP in the presence of *Rg*DAAO thus phenocopying the blocking action of d-AP5 (Fig. 3b). In conclusion, the excitatory PN-*to*-PV^+^ synapse can undergo homosynaptic eLTP that relies exclusively on the activation of postsynaptic NMDAR by d-serine.

**Figure 3.**
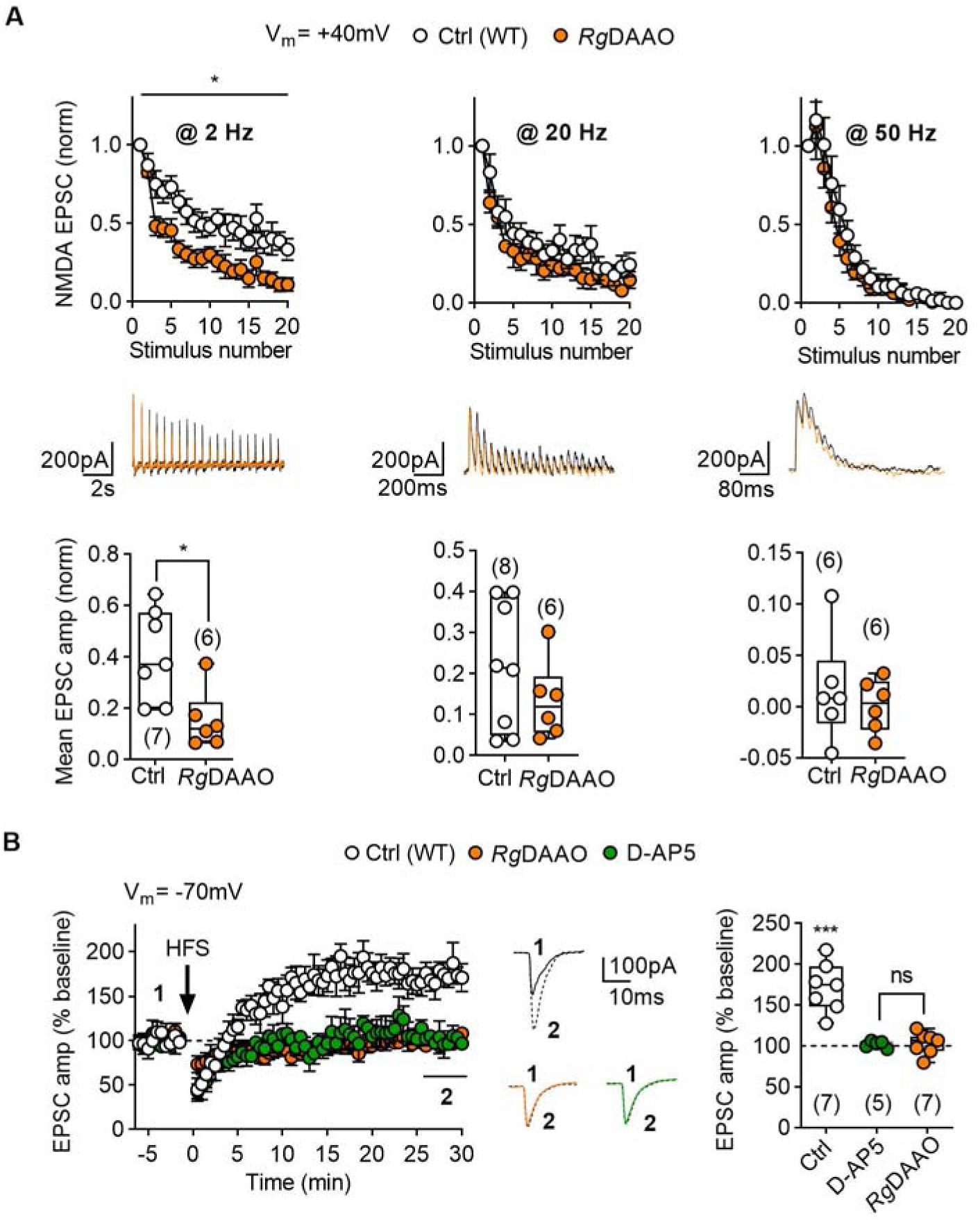
_D_-serine co-agonism of NMDARs is required for temporal summation and long-term plasticity at PN-*to*-FS-PV^+^ INs synapse. (**A**) Graphs show the normalized NMDA EPSCs peak amplitudes in response to stimulation trains applied at 2, 20 or 50 Hz in control (white) and when slices were treated with *Rg*DAAO (orange) to deplete d-serine. At 2 Hz *Rg*DAAO alters temporal summation resulting in a more pronounced attenuation (n= 7 Ctrl *vs* 6 *Rg*DAAO cells, *p*= 0.011, F(1,11)= 2.474, two-way ANOVA).). No difference was observed at either 20Hz (n= 8 Ctrl and 6 *Rg*DAAO cells, *p*= 0.1987, F(1,12)= 1.851, two-way ANOVA) or 50 Hz (n= 6 cells per condition, *p*= 0.450, F(1,10)= 0.619, two-way ANOVA) between control and *Rg*DAAO. Sample traces (middle) are shown comparing the Ctrl (black) and the *Rg*DAAO conditions. Bottom, boxplot graphs depict mean normalized NMDA EPSCs amplitude in Ctrl and *Rg*DAAO groups for the 5 last points at 2 Hz (*p*= 0.014, Mann-Whitney test), 20 Hz (*p*= 0.4948, Mann-Whitney test) and 50 Hz (*p*= 0.937, Mann-Whitney test). (**B**) Excitatory long-term potentiation (eLTP) at the PC-*to*-PV^+^ synapse is D-AP5 (green) sensitive. Left, time-course of LTP induced by 2×100 Hz tetanus stimulation (arrow) in control (white), D-AP5 or *Rg*DAAO group. Representative AMPA EPSCs are illustrated before (1, continuous lines) and after stimulation (2, dashed lines). Right, mean normalized amplitude of the 5 last minutes for each group indicated that LTP, normally induced in control (n= 7 cells, *p*= 0.016, Wilcoxon test *vs* 100), is absent both on D-AP5 (n= 4 cells, *p*= 0.4375, Wilcoxon test *vs* 100) and *Rg*DAAO (n= 7 cells, *p*= 0.4688, Wilcoxon test *vs* 100) conditions. \**p*< 0.05, \*\**p*< 0.01, \*\*\**p*< 0.001. All data are expressed as mean ± SEM.

### Glycine behaves as a spare co-agonist for synaptic NMDARs at FS-PV^+^ INs in a genetic mouse model of NMDAR hypofunction

Since d-serine is pivotal in regulating NMDARs, we next anticipated that the activity of NMDAR at FS-PV^+^ INs would be ultimately affected in the serine racemase knockout (SR^-/-^) mice (Fig. 4a), a translational model of NMDAR hypofunction caused by the loss of d-serine production (18–20, 59). These mice show no aberrant cortical lamination or visible loss of cell density (Fig. 4b) but the excitability of the mPFC FS-PV^+^ INs is markedly reduced in comparison to control wild-type (WT) littermate (Fig. 4c, Supplementary Fig. 3). Application of d-serine alone at a saturating concentration (20 µM) rescues both firing frequency and rheobase back to WT values (Fig. 4c). Unexpectedly, no significant differences were found in the voltage dependency and peak amplitude of NMDA EPSCs recorded from FS-PV^+^ INs in SR^-/-^ mPFC (Fig. 4d, e). In addition, NMDAR/AMPAR ratio was unchanged in SR^-/-^ slices when compared to WT slices (Fig. 4e) further confirming that under low electrical stimulation, synaptic parameters are globally preserved. We then reasoned that glycine, which is left intact in SR^-/-^ mice, could serve as a spare co-agonist to gate synaptic NMDARs. To test this hypothesis, we next examined the effect of incubating the SR^-/-^ slice with *Bs*GO to acutely reduce the ambient glycine. While *Bs*GO previously did not affect NMDA EPSCs or the excitability of FS-PV^+^ INs in WT slices (Fig. 2c,g), addition of *Bs*GO (0.2 U/mL) to SR^-/-^ slices remarkably reduced NMDA EPSCs by ∼60% without altering AMPA EPSCs, resulting in a net decrease of NMDAR/AMPAR ratio (Fig. 4e). These data demonstrate that glycine could serve as a spare co-agonist for gating synaptic NMDARs at excitatory synapses onto FS-PV^+^ INs when d-serine is chronically absent to control synaptic functions, but not firing activity.

**Figure 4.**
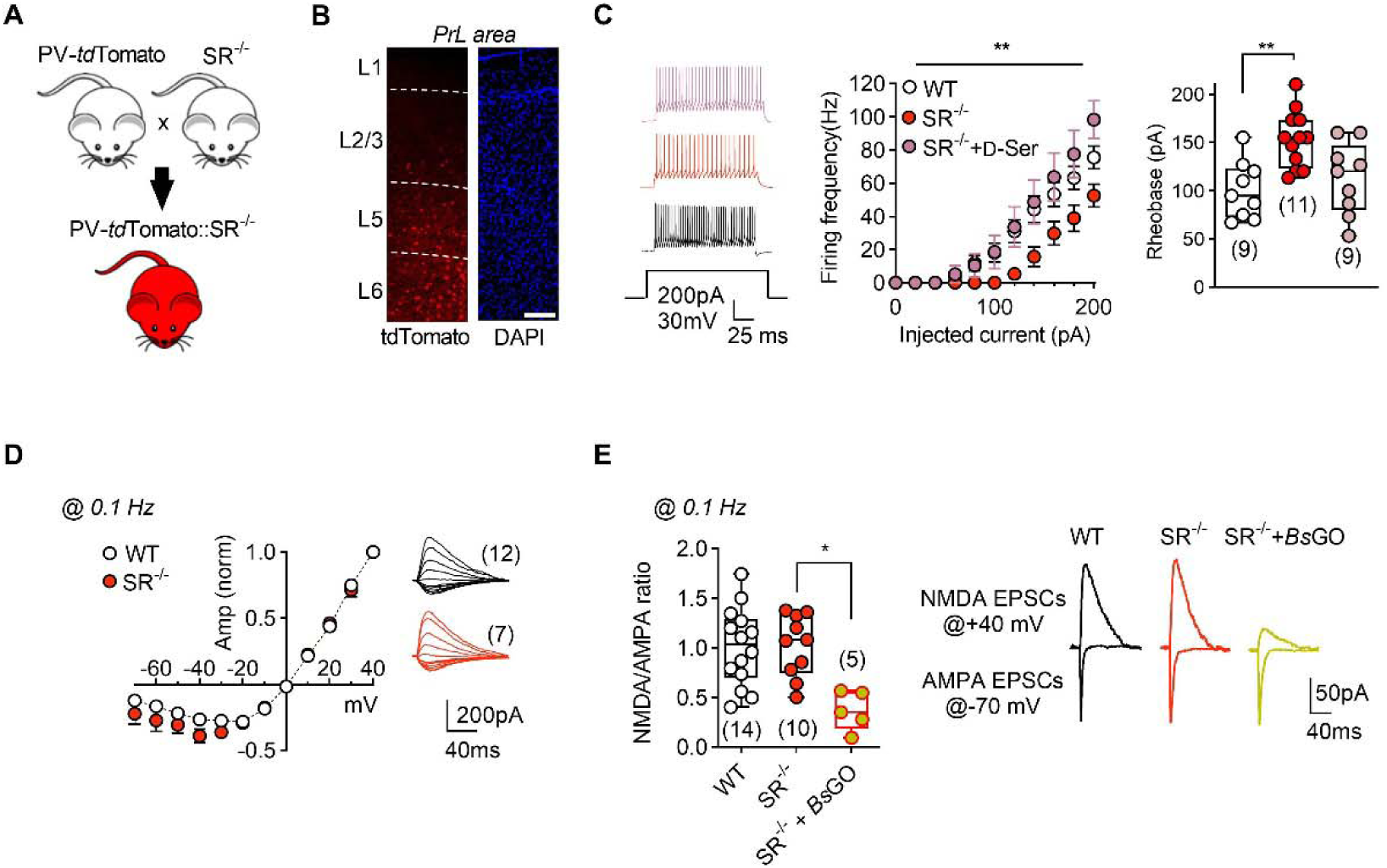
Glycine is a spare substitute of d-serine to regulate NMDARs at FS-PV^+^ interneurons. (**A**) PV^+^-*td*Tomato::SR^-/-^ mice were obtained by back-crossing PV^+^-*td*Tomato with SR^-/-^ mice. Only homozygous animals were used. (**B**) Representative confocal image of *td*Tomato expressing FS-PV^+^ INs in the mPFC of a SR^-/-^ mouse. DAPI staining reveals normal lamination and cell density in respect to WT mice. (**C**) Sample traces (left) showing the firing activity of FS-PV^+^ INs in response to a 200 pA current step injection recorded in WT (black trace), in SR^-/-^ mice before (red) or after bath application of d-serine (pink, 20 µM). Spike counts (middle) from graded current injections (0 to 200 pA in 20-pA steps) on interneurons for WT (n=9 cells), SR^-/-^ (n=11 cells, *p*=0.0004) and SR^-/-^ + d-serine (n=9 cells, *p*= 0.0003). Data were analyzed with two-way ANOVA and post-hoc Tukey’s test. Box plots (right) showing the associated effects on rheobase (SR^-/-^ *p*= 0.0072 *vs* Ctrl, *p*= 0.0917 *vs* SR^-/-^+ d-serine, Kruskal-Wallis test, post-hoc multiple comparison test). (**D**) I-V relationship of normalized NMDA EPSCs peak amplitude in SR^-/-^ (n= 7 cells) in respect to WT mice (n= 12 cells, *p*= 0.162, F(1,17)= 2.133, two-way ANOVA). (**E**) NMDA/AMPA EPSCs peak amplitude ratio recorded in WT (ie Ctrl, n= 14 cells), SR^-/-^ slices (n=10 cells, *p*> 0.9999 *vs* WT) and SR^-/-^ slices depleted of glycine with *Bs*GO (red-olive, n=5 cells, *p*= 0.014 *vs* SR^-/-^ or WT). Data were analyzed with Kruskal-Wallis test and post-hoc Dunn’s test. \**p*< 0.05, \*\**p*< 0.01, \*\*\**p*< 0.001. Data are expressed as mean ± SEM.

Noteworthy, NMDA EPSCs from FS-PV^+^ INs in SR^-/-^ mice systematically displayed a shorter rise time (Supplementary Fig. 6) but no change in decay kinetics, similar to what was observed upon acute ablation of d-serine with *Rg*DAAO. As rise time of EPSCs reflects the opening rate of the NMDAR channels, the mode of activation of the NMDARs in SR^-/-^ mice might likely differ from WT mice because of increased dwell time of neuromodulatory signals in the extracellular space, that would also explain the decrease in excitability of FS-PV^+^ INs in SR^-/-^ mice (Fig. 4c). To test this hypothesis, we first assessed the relative ambient level of co-agonist at FS-PV^+^ INs by recording the tonic current under a low concentration of the competitive NMDAR antagonist DCKA, which is selective for the co-agonist binding site. No significant difference was found in the action of DCKA on the holding current or synaptic noise between SR^-/-^ and control WT mice (supplementary Fig 9a-d). Therefore, the faster rise time of SR^-/-^ NMDA EPSCs seems not being a consequence of co-agonist spillover and changes in dwell time of the co-agonist in the extracellular space. Then, we wondered whether the levels of ambient glutamate may have changed upon genetic deletion of SR. To test this hypothesis, we employed the low-affinity and rapidly dissociating competitive AMPAR antagonist γ-DGG. At non-saturating concentrations, its potency is inversely depending on glutamate concentration (60). Inhibition of evoked AMPA EPSCs by bath applied 0.2 and 0.5 mM γ-DGG was lower in SR^-/-^ slices compared to WT (Supplementary Fig. 9e). These data indicate that the ambient level of glutamate is increased at synaptic sites in SR^−/−^ mice, suggesting that compensatory mechanisms for d-serine absence intervene, most likely explaining the changes in NMDA EPSCs kinetics and the loss of FS-PV^+^ INs excitability.

### Long-term synaptic plasticity of feedforward excitation is absent in mice lacking serine racemase

As glycine is sufficient to maintain basal activity of (at least a fraction of) synaptic NMDARs, we then investigated whether glycine could be recruited in SR^-/-^ mice when increasing the synaptic activity regime. Because at 50 Hz the synaptic attenuation and summation do not depend anymore on the degree of occupancy of the NMDAR co-agonist binding site (Fig. 3a), we thus restricted our evaluation of NMDA EPSCs at 2 and 20 Hz in SR^-/-^ mice. Interestingly, we found no difference with WT littermates at 2 Hz but revealed that rate tuning in SR^-/-^ slices was altered at 20 Hz (Fig. 5a). These results thus indicated that glycine could sustain NMDAR activity in SR^-/-^ mice at low firing rate (e.g θ regime) but beyond, the system became invariably unstable in the chronic absence of d-serine. Indeed, while high-frequency stimulation consistently induced robust NMDAR-dependent LTP in WT mice, this protocol was ineffective in triggering eLTP in SR^-/-^ mice (Fig. 5b). The absence of eLTP was not associated with changes in presynaptic parameters as PPR in WT and SR^-/-^ mice was similar (PPR∼1.5) before and after LTP (Supplementary Fig. 8). Therefore, glycine contrary to d-serine, and despite its action on NMDARs at low synaptic regime, is not recruited in the formation of eLTP at PN-*to*-PV^+^ synapses. Overall, these results argue that glycine could compensate for the chronic absence of d-serine (e.g., SR^-/-^) to support the demand of NMDAR activity in FS-PV^+^ INs at low but not at high synaptic activity regimes, in striking contrast to data gathered on synapses between PNs, where no evident deficit were observed in the d-serine deficient mice (39, 47, 61, 62).

**Figure 5.**
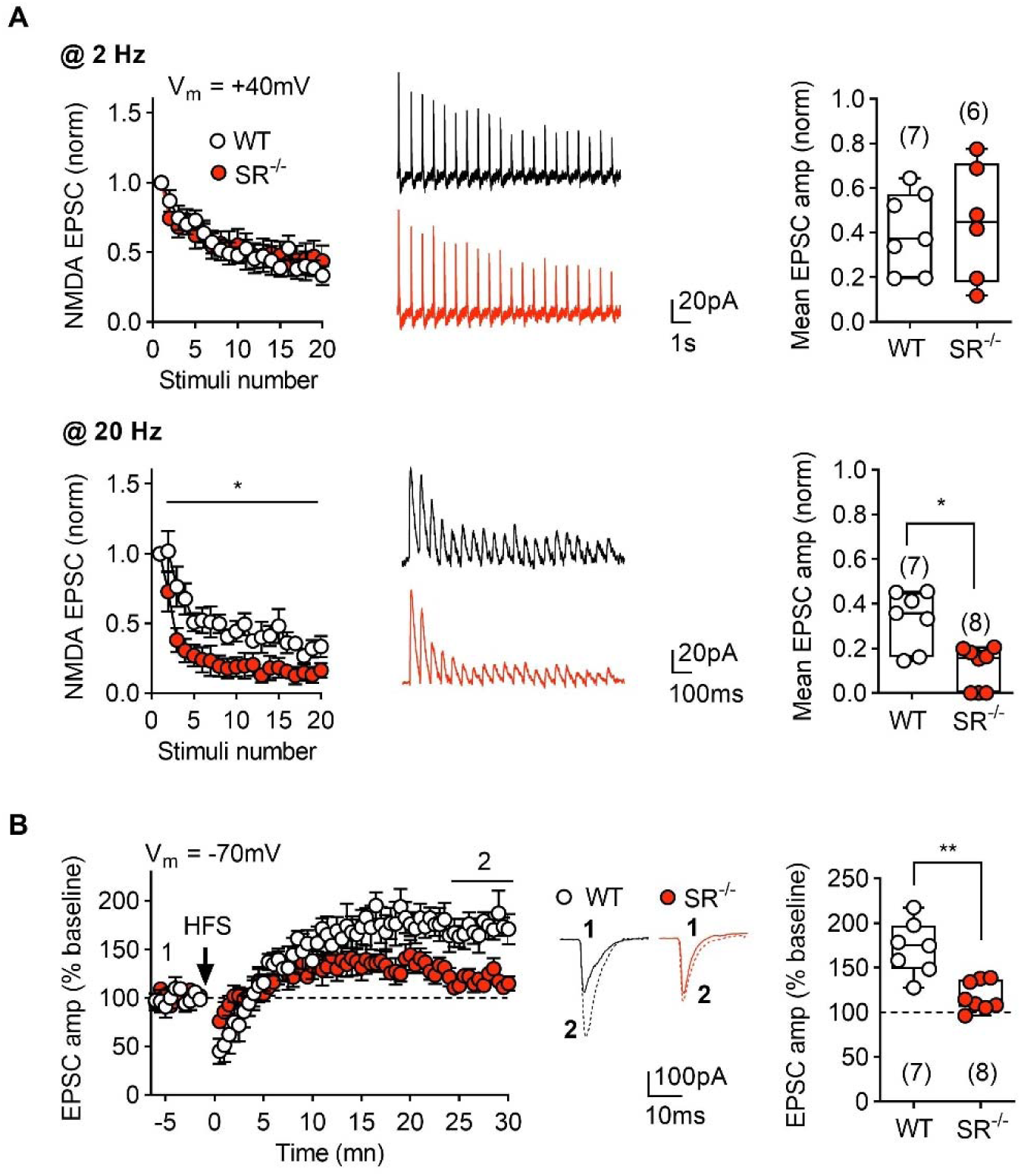
**Temporal summation and synaptic plasticity of excitatory PN-*to*-PV^+^ synapse is impaired in SR**^-/-^ **mice. (A)** At 2 Hz, temporal summation in SR^-/-^ (red) FS-PV^+^ INs was normal compared to WT (white, n= 7 WT and 6 SR^-/-^ cells, *p*= 0.920, F(1,11)= 0.011, two-way ANOVA). In contrast, at 20 Hz, chronic deficiency in d-serine impaired temporal summation (n= 7 WT and 8 SR^-/-^, *p*= 0.020, F(1,13)= 7.047, two-way ANOVA) in line with the effect of *Rg*DAAO (see Fig 5, SR^-/-^ vs *Rg*DAAO *p*> 0.9999, post-hoc Bonferroni’s test after two-way ANOVA). Comparison of the mean normalized NMDA EPSCs amplitude of the 5 last points at 2 and 20 Hz for SR^-/-^ and WT groups reveals no difference at 2 Hz (*p*= 0.945, Mann-Whitney test). On the contrary, mean amplitude at 20 Hz is significantly reduced for SR^-/-^ (*p*= 0.028, Mann-Whitney test). **(B)** HFS-induced excitatory long-term potentiation (eLTP) in SR^-/-^ mice showed a significant reduction in comparison to WT (n= 7 WT and 8 SR*^-/-^* cells, *p*= 0.002, Mann-Whitney test). However, the short-term potentiation 10 min post stimulation was observed in both groups. \**p*< 0.05, \*\**p*< 0.01. Data are expressed as mean ± SEM.

### _D_-Serine but not glycine regulates FS-PV^+^ INs-mediated feedback inhibition

According to the NMDAR hypofunction hypothesis, loss of NMDAR function at FS-PV^+^-INs would hence result ultimately in decreasing output GABAergic neurotransmission. Thus, we next explored whether and how the NMDARs co-agonist site occupancy at FS-PV^+^ INs may regulate the strength of PV^+^-*to*-PN inhibitory synapse in the mPFC. In all experiments, to enable the sole manipulation of NMDARs at the presynaptic cell without interfering with NMDARs at PNs, MK801 (2 mM) was applied to the postsynaptic PN *via* the patch recording pipette (Figure 6a,b), thus acutely blocking NMDARs in the recorded excitatory PN. We found that NMDA application (10 µM) increased the action potential (AP)-dependent TTX-sensitive spontaneous inhibitory postsynaptic currents (sIPSCs) frequency but not their amplitude (supplementary Fig 10) while bath applied d-AP5 (50 µM) decreased sIPSCs frequency. Interestingly, the frequency of sIPSCs which was depressed in SR^-/-^ mice could be restored to normality upon application of d-serine (Supplementary Fig. 10). Since NMDARs are absent from axonal terminal of layer 5 PV^+^-INs in the mPFC (Prof Z Nusser, personal communication), we reasoned that NMDARs activation might enhance GABA release *via* subthreshold depolarization of the FS-PV^+^ IN somatodendritic arbor that propagates to the axon terminals. Bath-applied NMDA potentiated the perisomatic electrically elicited IPSCs while d-AP5 had the opposite effect (Fig 6c, supplementary Fig. 11). Such elicited IPSCs did not involve perisomatic GABA synapses from cholecystokinin-positive (CCK^+^) INs since ω-conotoxin, which blocks GABA release from CCK^+^ INs (31), did not affect the IPSCs peak amplitude nor prevented the potentiating effect of NMDA (Supplementary Fig. 12). On the contrary ω-agatoxin IVB, which blocks the release of GABA by FS-PV^+^ INs, nulled the recorded IPSCs and prevented the potentiating effect of bath applied NMDA and nulled inhibitory LTP (LTP, Supplementary Fig. 12).

**Figure 6.**
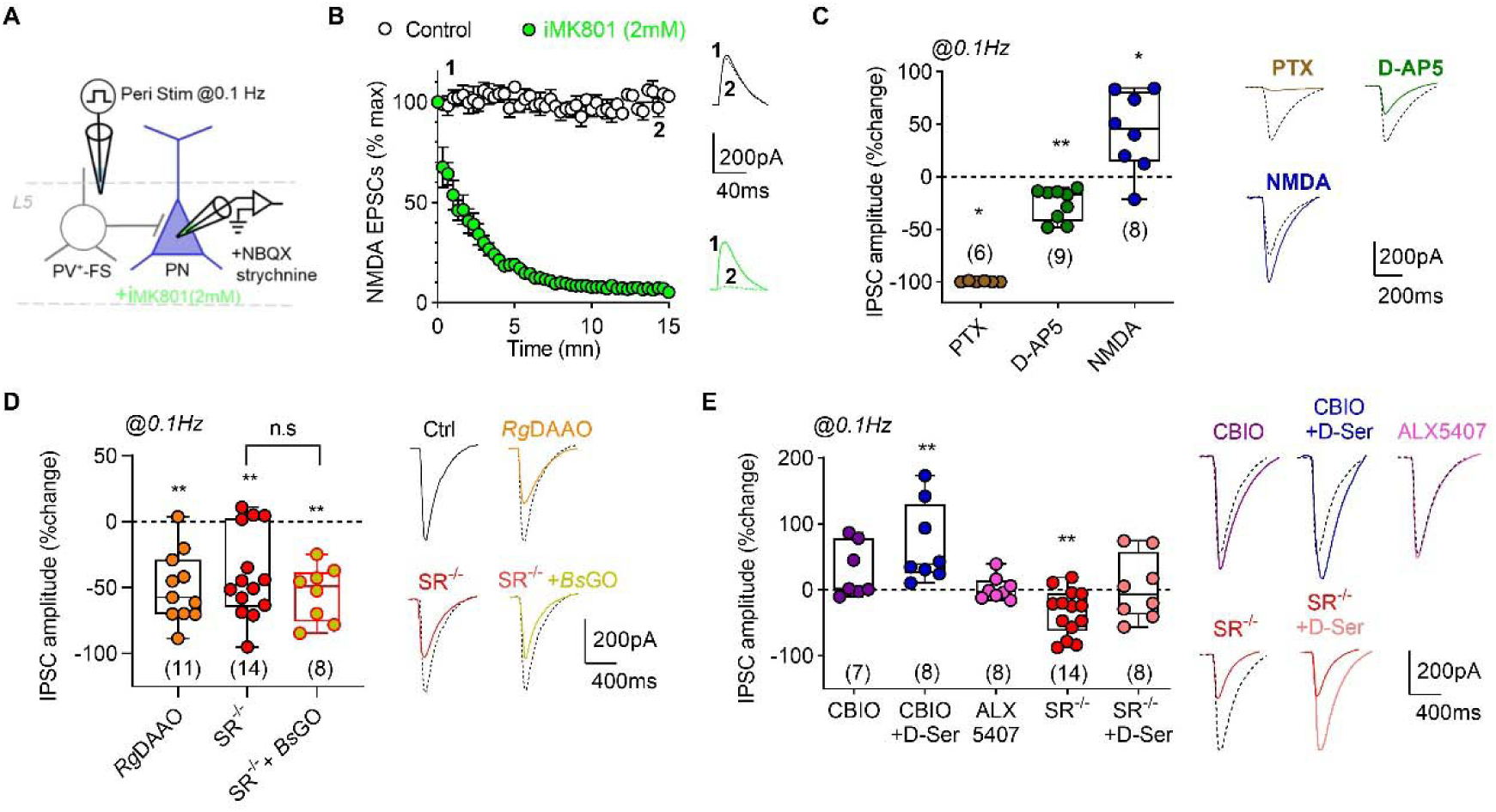
NMDARs co-agonism by d-serine controls FS-PV^+^ interneurons mediated feedback inhibition to pyramidal neurons. **(A)** Schematic diagram showing the position of the recording electrode in mPFC layer 5, where electrically evoked synaptic GABA_A_ currents from pyramidal neurons (PN) were recorded in WT or SR^-/-^ mice, with inclusion of MK801 in the patch pipette for isolation of NMDAR contribution solely from FS-PV^+^ INs. **(B)** Time course (left) of the effective inhibitory action of MK801 (light green) when included in the patch pipette in NMDA EPSCs peak amplitude. Sample traces (right) of NMDA EPSCs recorded at +40 mV are represented before (1) and after 15 min action of MK801 or vehicle (2). **(C)** Effect of NMDAR modulation on electrically evoked PTX-sensitive IPSCs. While D-AP5 (dark green, n= 9 cells) partially reduced IPSCs, NMDA (dark blue, n= 8 cells) had the opposite action. PTX (brown, n= 6 cells) nulled IPSCs. Superimposed example traces (dashed line, Ctrl, signed rank Wilcoxon test; PTX: *p*= 0.0313; D-AP5: *p*= 0.0039; NMDA: *p*= 0.0391 *vs* Ctrl). **(D)** Effect of modulating the co-agonist site occupancy of NMDARs on IPSCs. Box plots (left) show the percentage of changes induced by *Rg*DAAO (orange, n= 11 cells) and in SR^-/-^ (red, n= 14 cells) mice in comparison to control (white, n= 11 cells). Note that *Rg*DAAO in WT caused a similar decrease as in SR^-/-^ mice. Application of the glycine scavenger *Bs*GO (olive, n= 8 cells) to SR^-/-^ slices did not further decrease the observed amplitude (*Rg*DAAO: *p*= 0.002; SR^-/-^: *p*= 0.005; SR^-/-^+*Bs*GO: *p*= 0.0078 all *vs* Ctrl, Wilcoxon signed rank test). Examples traces (right) are shown. Traces are shifted for easy comparison (dashed line, Ctrl). **(E)** IPSCs responses to pharmacological modulation of glycine and d-serine levels. Potentiating d-serine co-agonism with CBIO (purple, n= 7 cells) or CBIO + d-serine itself (dark blue, n= 8 cells, *p*= 0.0078, Wilcoxon signed ranked test) increased IPSCs, while potentiation of glycine co-agonism with ALX5407 (pink, n= 8 cells) did not. Reduction of IPSCs amplitude in SR^-/-^ mice (n= 14 cells, *p*=0.0023 Wilcoxon signed ranked test) was reversed by d-serine (20 µM) (light red, n= 8 cells, *p*>0.9999, Wilcoxon signed ranked test). \**p*< 0.05, \*\**p*< 0.01. Data are expressed as mean ± SEM.

Then, we explored the relative roles of NMDARs co-agonists in regulating the PV^+^-*to* PN synaptic strength. *Rg*DAAO (0.2 U/mL), which acutely depletes d-serine, depressed the evoked IPSCs peak amplitude and a similar reduction was observed when recordings were performed in slices from SR^-/-^ mice (Fig. 6d-e, supplementary Fig 11). In strong contrast with the PN-*to*-PV^+^ synapse (Fig. 4e), *Bs*GO had no additional inhibitory action on evoked IPSCs recorded in SR^-/-^ mice (Fig. 6d) excluding a compensatory role for glycine at this synapse. Likewise, ALX5407 failed to increase the evoked IPSCs in WT slices (Fig 6e). On the contrary, application of CBIO alone or in combination with d-serine increased evoked IPSCs and bath applied d-serine restored the depressed IPSCs to control value in SR^-/-^ mice (Fig. 6e, supplementary Fig 11). Altogether these data indicate that d-serine, but not glycine, is able to control the PV^+^-*to*-PN synaptic strength by modulating the somatodendritic NMDARs.

### Glycine supports short term and long-term synaptic plasticity of FS-PV^+^ INs-mediated feedback inhibition in SR knock-out mice

Finally, we investigated the contribution of NMDARs co-agonism by d-serine on temporal summation and long-term synaptic plasticity. Accordingly, and as previously done for the PN-*to*-PV^+^ excitatory synapse, the PV^+^-*to*-PN inhibitory synapse was subjected to different trains of synaptic stimulation. At 2Hz and 50Hz, the temporal summation of evoked IPSCs was not affected by acute d-serine depletion with *Rg*DAAO (Fig. 7a). Conversely, short-term depression was dampened at 20 Hz (Fig. 7a) indicating that d-serine is necessary to fully express synaptic fatigue of GABAergic outputs. Finally, we probe for the role of d-serine in inhibitory LTP (iLTP). No iLTP was induced using a hebbian HFS protocol used previously at the excitatory synapse. On the contrary readable iLTP could be induced following TBS stimulation in WT slices (Fig. 7b) with the postsynaptic cell maintained at −70 mV. This non-hebbian type of LTP, which depends on dendritic NMDARs (supplementary Fig 11) as also prevented by d-AP5, was maintained in SR^-/-^ mice (Fig. 7b). Together, these data support the idea that glycine, which is left intact in these conditions, is recruited at the PV^+^-*to*-PN inhibitory synapse during increased synaptic regime to support rate tuning and long-term synaptic plasticity.

**Figure 7.**
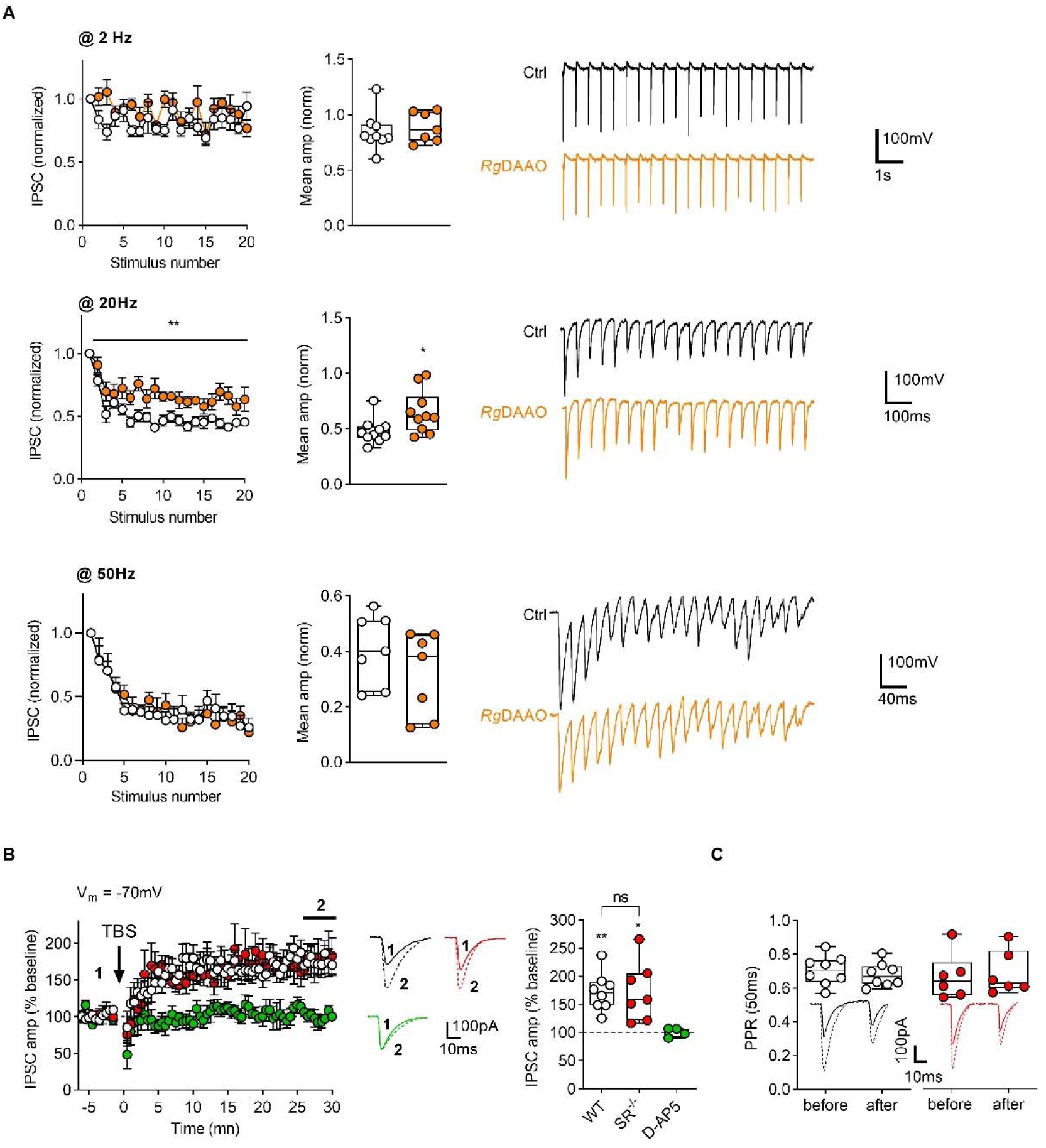
_D_-serine co-agonism of NMDARs is not required for temporal summation and long-term plasticity of inhibitory PV^+^-*to*-PN synapses. **(A)** Normalized IPSCs amplitudes showed graded attenuation in response to stimulation trains applied at some frequencies. Top, at 2 Hz, acute d-serine depletion with *Rg*DAAO (orange) did not affect temporal summation (n= 9 control and 7 *Rg*DAAO cells, *p*= 0.0926, F(1,14)= 3.258, two-way ANOVA). Middle, at 20 Hz, *Rg*DAAO had a small but significant inhibitory effect (n = 9 control and 10 *Rg*DAAO cells, *p*= 0.0010, F(1,14)= 17.05, two-way ANOVA). Bottom, at 50 Hz, no difference was observed between control (n= 7 cells) and *Rg*DAAO (n= 7 cells) groups (*p*= 0.9993, F(1,12)= 7,558.10^-7^, two-way ANOVA). Boxplot graphs depict mean normalized IPSCs amplitude in Ctrl and *Rg*DAAO for the 5 last points at 2 Hz (*p*= 0.6806, Mann-Whitney test), 20 Hz (*p*= 0.0288, Mann-Whitney test) and 50 Hz (*p*= 0.2593 Mann-Whitney test). (**B**) Inhibitory long-term potentiation (iLTP) at the PV^+^-to-PN synapse is also D-AP5 (green) sensitive but independent to d-serine co-agonism. Left, time course of iLTP induced by a theta burst stimulation protocol (4-stimuli at 100Hz repeated 25 times at 5Hz) in control, D-AP5 or SR^-/-^ (red) group. Representative IPSCs are illustrated before (1, continuous lines) and after stimulation (2, dashed lines). Right, mean normalized amplitude of the 5 last minutes for each group indicated that LTP is normally induced in control (n= 8 cells, *p*= 0.008, Wilcoxon test *vs* 100) and SR^-/-^ (n= 7 cells, *p*= 0.016, Wilcoxon test *vs* 100) but is absent in D-AP5 (n= 4 cells, *p*> 0.999, Wilcoxon test *vs* 100) condition. (**C**) PPR analysis before and after iLTP in WT and SR^-/-^ mice. Box plots show no variations in the PPR between groups. \**p*< 0.05, \*\**p*< 0.01. Data are expressed as mean ± SEM.

## Discussion

FS-PV^+^-INs are critical cellular elements regulating cortical functions which are disrupted in psychiatric neurodevelopmental disorders (1–10), hence NMDARs populating them are considered as primary therapeutical targets (63). We carried out a quantitative analysis to explore the physiological conditions that specify the actions of d-serine and glycine in controlling the activity and functions of NMDARs at FS-PV^+^-INs. We nowreport that functional synaptic NMDARs are substantially detectable in mPFC FS-PV^+^-INs beyond the neonatal window as classically reported. By combining selective acute pharmacological and genetic manipulations, we reveal that glycine and d-serine have complementary actions in regulating somatodendritic NMDARs functions at FS-PV^+^-INs in the adolescent mPFC. One key conclusion from the present study is that d-serine but not glycine modulates the excitatory drive of cortical GABAergic PV^+^-INs by acting on NMDARs. We demonstrate that acute or chronic depletion of d-serine induces NMDARs hypofunction impairing burst firing, with alterations in excitatory short-term and long-term plasticity at PN *to* FS-PV^+^-IN synaptic connections. On the contrary at FS-PV^+^-IN *to* PN synapses, we report a synaptic activity-dependent shift in the identity of the co-agonist from d-serine to glycine supporting normal inhibitory summation and long-term potentiation in SR^-/-^ mice where glycine can fully serve as a substitute when high-frequency activity is required.

PV^+^-INs represent distinct populations of neurons with distinct biophysical, morphological, and thus with different cortical functions. Here, inhibitory neurons were identified using PV^+^-tdTomato reporter mice and therefore no distinction between the chandelier and basket cells types neurons could be easily made at first. However, we are confident that we only selected the basket cells in our study. Indeed, even if both cell populations are present in the mouse mPFC, only the basket cells are found in deep cortical layers (64). Furthermore, we systematically include in our recordings only the PV^+^ neurons with fast-spiking discharges and no attenuation which are typical biophysical features of the basket cells. This electrophysiological trait refers to the short kinetics of the action potential (Supplementary Fig. 1) consequential of the selective expression of Kv3.1b potassium channels in those cells with no overlap with others GABAergic or excitatory neurons.

_D_-Serine and glycine can evenly bind to the glutamate insensitive diheteromeric GluN1-GluN3 and to the homotetrameric GluD receptors (12, 13) that also populate excitatory and inhibitory neurons. However, the modulatory actions of the co-agonists at FS-PV^+^-INs we are now reporting most likely do not engage these unconventional glutamate receptors since all elicited NMDAR-dependent responses were buffered by the non-competitive blockers D-AP5 or MK801. Furthermore, analysis of the current-voltage (I-V) curves of NMDA-EPSCs recorded at FS-PV^+^ INs shows voltage-dependent rectifications typical of GluN1-GluN2 containing receptors in physiological Mg^2+^ (1.5 mM). By using fluorescent *in situ* hybridization, we revealed that GluN2B and GluN2/D are the two major subunits expressed by mPFC PV^+^-INs aligning with the literature (32–36, 65). GluN3A transcripts are also present albeit at a relatively low level (supplementary Fig. 4). Although, we cannot exclude that GluN3A-GluN1 could segregate to form a separate pool of diheteromeric NMDARs, most likely GluN3A enter in the composition of triheteromeric NMDARs as suggested by the total blockade of the responses we are monitoring in the presence of MK801 or D-AP5. By using selective pharmacology, we found that both GluN2B and GluN2C/D contribute to synaptic NMDAR currents in young adults. The presence of GluN2B, GluN2D and to some extend of GluN3A at FS-PV^+^ INs in the mPFC well aligns with the recognized protracted development of cortical FS-PV^+^ INs and the PFC compared to other interneurons and brain areas (22–24). We have not explored the presence of GluN2A. However, the presence of GluN2A in FS-PV^+^ INs is unlikely despite the fact that INs in layer 5 have been found to be sensitive to GluN2A antagonism (66). The long deactivation time course of our NMDA-EPSCs suggest that GluN2A is absent as the integration of GluN2A in NMDARs assemblies would be expected to shorten the decay regardless of the presence of others subunits. Furthermore, no obvious remodeling in the subunit composition of NMDARs seems to occur in the SR null mutant mice as the peak amplitude and the decay time of the NMDA EPSCs were not altered (supplementary figure 6) suggesting that the postsynaptic proteome of FS-PV^+^ INs is conserved. This is remarkable since SR has been found to interact with several NMDAR- and AMPAR-interacting postsynaptic proteins including PSD95 (67, 68) in cortical neurons. Deletion of SR would be expected to affect the development and stability of glutamatergic synapses (68). Therefore, glycine could fully substitute to d-serine in maintaining the integrity of the glutamatergic postsynapse. SR^-/-^ mice do not show any anatomical alteration in the layering of the cerebral wall and no deficiency in the density of PV^+^-tdTomato cells (Figure 4), suggesting that a decrease in GABAergic INs and notably of FS-PV^+^ cells under persistent absence of d-serine is unlikely (21). Furthermore, we found that the occupancy of the co-agonist site of the synaptic NMDARs controlling the excitatory input is not affected in SR^-/-^ mice (Figure 4) which also opposes to a reduced number of GABAergic cells. Therefore, we can conclude that the alterations in inhibitory/excitatory balance, brain oscillations and behavior observed in SR^-/-^ mice (18–20, 47, 61, 62) are not reflected by any macroscopic anatomical defects during development at least in the PFC.

Subcellular analysis using quantitative high-resolutive electron microscopy to map the membrane expression of GluN1 revealed that NMDARs are abundantly present at the excitatory postsynaptic density of PN-to-PV connections while they are absent from Kv3.1b^+^ axon terminals (Zoltan Nusser, personal communication). Therefore, NMDARs are strictly confined to the somatodendritic compartments of adolescent FS-PV^+^ INs where they are controlling the excitatory input and the GABAergic output inhibition we are here reporting. This observation raises an intriguing scenario where the apparent same populations of receptors gated by either d-serine or glycine would exert different roles on the input and output functions of FS-PV^+^ INs in an activity-dependent manner.

We show that the identity of the co-agonist (eg d-serine) at GABAergic neurons is not dictated by the level of afferent excitatory inputs activity and that glycine can only sparely replace d-serine. Indeed, compensation by glycine of NMDAR gating in SR^-/-^ mice or under acute loss-of function of d-serine (e.g *Rg*DAAO) is incomplete and deficits of excitatory drive of PV^+^ interneurons already appear at low frequency range (∼2-20 Hz). Since glutamate is fully saturating NMDARs during synaptic transmission, d-serine availability is the limiting factor for optimal NMDAR activation at PV^+^ interneurons. Accordingly, despite normal synaptic transmission at low synaptic regimes (<2Hz), HFS-induced excitatory LTP is occluded in SR^-/-^ mice just phenocopying the effect of *Rg*DAAO treatment. Yet, iLTP is maintained in SR^-/-^ mice suggesting that glycine could fully substitute to d-serine in maintaining plasticity of inhibitory output. This likely provide a mechanism to explain the impaired E/I balance and inefficient information processing reported in the SR^-/-^ mice

The mechanisms underlying the selective action of d-serine over glycine at FS-PV^+^ inhibitory neurons are not evident. Although the NMDARs subunits composition may differ between excitatory and inhibitory neurons (12, 13), both neuromodulators display nearly similar sensitivity to NMDARs independently of their subunits composition (12, 13). The partial compensatory action of glycine at GABAergic neurons in SR^-/-^ mice may rather reflect differences in the dynamics and availability of the co-agonists between excitatory synapses onto PNs *vs* inhibitory neurons, as already observed for glutamate (60). By expressing the GlyT1, astrocytes are critically involved in regulating the ambient levels of glycine at the proximity of excitatory neurons (38–40). Hence, the synaptic microenvironment of GABAergic and glutamatergic neurons would be largely influenced by the morphological interposition of individual synapses with astrocytes (69, 70). Excitatory neurons establish close contact with astrocytes leaflets while GABAergic neurons notably the PV^+^-INs establish sparse contact with glia (70) although still contributing to inhibitory synaptic signalization. In line with these observations, pharmacological blockade of GlyT1 by ALX5407 to build up ambient glycine levels increased synaptic NMDAR currents when IPSCs remained largely unaffected. Unlikely, inhibition of DAAO by CBIO to boost extracellular d-serine availability increased firing activity of PV^+^-INs and their input and output synaptic couplings with PNs. We did not address precisely the participation of other circuits elements or astrocytes in the neuromodulation induced by d-serine and/or glycine. However, contribution of other cellular elements outside FS-PV^+^ INs and PNs is negligible since networks activity was down regulated in our experiments. However, DAAO is found in neurons and in non-neuronal cell types and it is possible that the effect of CBIO could be the result of a complex circuit phenomena involving multiple and distinct cellular groups. Furthermore, our observation that the effect of GlyT1 with ALX5407 or DAAO with CBIO is systematically greater on NMDA-EPSCs amplitude when they are minimal (CBIO) or absent (ALX5407) on the IPSCs, also support the idea that d-serine and glycine in the synaptic cleft needs to reach subthreshold levels to efficiently induce depolarization of the FS-PV^+^-IN to modulate the output inhibition. The presence of active DAAO in the forebrain has been highly questioned for decades and therefore its role in d-serine catabolism remains an opened issue (43–46). Our previous work indicated that the potentiating effect of CBIO on excitatory synaptic transmission in the hippocampus circuitry was nulled in SR^-/-^ mice thus demonstrating that the effect of the compound on inhibiting DAAO was caused by an elevation in d-serine (40). Here, our pharmacological experiments using CBIO further evidence that DAAO is well present in the forebrain and is the enzyme catabolizing d-serine but most importantly, show that DAAO by controlling d-serine disposition plays a critical role in controlling the activity of FS-PV^+^ INs, alongside with their excitatory and inhibitory synaptic connections. We thus confirm the positioning of DAAO inhibitors as a promising pharmacological approach to boost cognitive performance through the up state of NMDARs at GABAergic neurons (71).

The most parsimonious interpretation for the difference in glycine and d-serine gating NMDARs is to propose the existence of at least two populations of NMDARs that are differently distributed along the somatodendritic arbor of the FS-PV^+^ INs. These different pools of NMDARs would be accessible for gating by d-serine and/or by glycine depending on the physiological needs. Accordingly, the d-serine sensitive NMDARs at PV^+^-INs would be preferentially located at proximal dendrites or on the soma where they are prompt following activation to initiate strong depolarization of the FS-PV^+^ INs. On the contrary, activation of more distal dendritic NMDARs by glycine would produce only a weak subthreshold depolarization of the soma because distal dendritic spikes are subject to significant attenuation by dendritic cable properties (72,73), and therefore would require higher elevated network activity to induce firing of FS-PV^+^ INs. Aligning with this scenario, we found that glycine is recruited during stronger depolarization or increased synaptic activity regime to control cell firing or inhibitory synaptic plasticity under persistent depletion of d-serine in SR^-/-^ mice. Distally-generated dendritic activation is known to play a critical role in inhibitory synaptic plasticity and complex behavior and would explain why SR^-/-^ mice display E/I imbalance and impaired EEG frequency spectra but only partial cognitive and synaptic plasticity deficits. Still, this scenario does not explain why when increasing the synaptic activity regime, glycine is not recruited to induce excitatory synaptic plasticity in the absence of d-serine (Figs 3 and 5). How can we reconcile the importance of the distal inputs triggered by glycine with the necessity of somatic excitation generated by d-serine? As already reported for Layer 5 pyramidal neurons in the hippocampus and neocortex (72, 73), it is plausible that somatic depolarization induced by excitatory synaptic input (under d-serine control) can be used to amplify the response at the soma to a subthreshold dendritic depolarization induced by the glycine sensitive NMDARs. Accordingly, d-serine by gating somatic NMDARs would act as a permissive factor facilitating the induction of plasticity driven by the distal synaptic inputs triggered by glycine. Aligning with this model, blocking the somatodendritic voltage-gated Na^+^ channels activation with QX314 did not prevent the formation of eLTP which relies on d-serine but not on glycine-dependent gating of NMDARs.

By reporting that pharmacological or genetic-induced down regulation of d-serine levels did alter the firing activities of the PV^+^ cells in a NMDAR-dependent manner and basal inhibitory synaptic transmission, we demonstrate that d-serine is the major permissive factor engaged in the tonic control of NMDAR in physiological conditions. A role for glycine in wild-type mice (e.g when d-serine is present) was unmasked only when measuring NMDA-EPSCs after blocking the GlyT1 with ALX5407 (Fig. 2) or only when strong depolarization was used to induce firing activity of PV^+^ cells or in controlling inhibitory plasticity (Fig. 7). This apparent discrepancy could be related to the fact that firing activities are recorded at the resting membrane potential under current clamp thus better preserving the physiological conditions. NMDAR synaptic currents were collected at + 40 mV to relieve the potential-dependent Mg^2+^ block of the channel and in response to non-minimal perisomatic stimulations. These experimental conditions are prompted to place the recorded cell and its local environment in an upper depolarizing state that may have unsilenced some NMDARs normally quiescent at physiological membrane potentials and thus may favor the recruitment of glycine. Aligning with this hypothesis, we found that *Bs*GO could effectively attenuate the firing of PV^+^-INs when injected currents are > 250 pA to depolarize the cells. Yet, during sharp wave ripples, high-frequency oscillations associated with memory formation corresponding to epochs of elevated cortical activity, PV^+^-INs exhibit prolonged depolarizations to voltages at which NMDAR Mg^2+^ block is substantially reduced. Furthermore, this scenario is also consistent with the idea that activation of extrasynaptic NMDARs by either d-serine or glycine contribute to GABAergic interneuron excitability in the mPFC and to behavioral performances (74). Burst firing of PV^+^-INs promote the reliability of somatic inhibitory drive to PNs thus playing a cardinal role in the temporal precision of local pyramidal firing and in the generation of coherent oscillations enabling the synchronization of large PN populations and proper behaviors (1–10). Recent findings suggest that NMDARs in PV^+^-INs enhances the probability of GABA release in the mPFC (31). In the circuit context, bursts of excitatory input onto PV-INs driven by glycine and/or d-serine would enable fine-tuning of their firing and their basal inhibition of PNs but also the adaptation of the temporal contrast of their responses to subsequent synaptic inputs, thus modulating feedforward inhibition on demand. We found that the absence of d-serine results in impaired firing activity in SR^-/-^ mice concurrently to alterations in inhibitory and excitatory short-term plasticity at 20Hz which indicates that disinhibition is increased under these conditions. In parallel, we also showed that the persistent depletion of d-serine leaves FS-PV^+^ INs in a more depolarized state relative to WT IN types (Supplementary Fig 3) making them more quiescent to the activation by low excitatory synaptic inputs by lowering the driving force for glutamate-induced currents. We reason that this likely results in a loss of their potency to act as feedforward inhibitory elements. Interruptions of PV^+^ INs firing are physiological safe metabolic events that preserve neurons from fatigue and provide a necessary disinhibitory circuit mechanism favoring spike generation in pyramidal cells. Inhibition of PV^+^ INs supports the temporal contrast necessary for learning and memory (75, 76). During complex behavioral tasks and during spatial and social exploration, there is a continual bombardment of FS-PV^+^ INs by excitatory synaptic inputs. As FS-PV^+^ INs in SR^-/-^ mice are more depolarized then less sensitive to excitatory inputs, they will be also less prone to these silent periods. We reason that the non-physiological persistent down state of FS-PV^+^ INs in SR^-/-^ mice and disinhibition of PN would recapitulate the symptoms found in these mice.

Persistent high-frequency activity of PFC neurons and coordinated network activity is required under normal conditions to support on-going sensory information processing which is disrupted in many neuropsychiatric disorders including SCZ. This would require notably the ability for FS-PV^+^ INs to mount and maintain high synaptic activity regimes. Acute or persistent depletion of d-serine results in dose-dependent disruptions of the excitatory drive of FS-PV^+^ INs to sustain high-frequency synaptic transmission (Supplementary Fig 12). Yet, we show that ambient level of glutamate is increased in SR^-/-^ mice relative to WT mice while the ambient levels of the co-agonist (ie glycine) remain unaffected. This could reflect either a dysfunction of reuptake glutamate systems notably by astrocytes or more likely an increased probability of glutamate release under persistent depletion of d-serine. Additional studies are warranted to examine the potential alterations of synaptic vesicles recycling properties in glutamatergic neurons and FS-PV^+^ INs under NMDAR hypofunction.

Collectively, our observations offer a mechanistic explanation why SR^-/-^ mice show impairments in gamma oscillations and social interactions (59, 60). In addition to unearth NMDAR hypofunction at PV^+^ interneurons in SR^-/-^ mice, our results offer a molecular and circuit-based explanation for how NMDAR hypofunction translate in defective FS-PV^+^ INs functions. We further highlight that the level of occupancy of the NMDAR co-agonist binding site depends on the cell type reflecting the divergent functional and most likely morphological properties of excitatory contacts onto PNs relative to interneurons (69, 70). Indeed, at PN-to PN excitatory synapses, it has been shown that both glycine and d-serine cooperate to regulate NMDAR-dependent synaptic plasticity (38–40, 62). Under pathological conditions of chronic d-serine absence as occurring in the SR^-/-^ mice, glycine can fully replace d-serine to support excitatory synaptic plasticity (39, 61, 62) especially in the mPFC (47) which is not the case for excitatory synapse onto FS-PV^+^-INs as described here.

Lastly, besides shifting our vision of brain circuit’s physiology, our study has translational relevance by fully justifying current therapeutics strategies targeting the co-agonist site of NMDARs by d-serine in the management of SCZ and others neurodevelopmental brain disorders linked to NMDAR hypofunction (11). Indeed, NMDAR hypofunction at PV^+^-INs has been invariably proposed to play a pivotal role in the pathogenesis of many psychiatric disorders, such as SCZ (5, 11) and studies suggest that these neurons should be considered as primary targets to more efficient antipsychotics. Our study connects these different emerging concepts by reporting that loss-of-function of d-serine induces selective NMDAR hypofunction at FS-PV^+^-INs, and that DAAO inhibition (i.e with CBIO) increases firing activity of FS-PV^+^-INs but not of PNs, and by doing so potentiate the excitatory input and the inhibitory tone. Although FS-PV^+^-INs may be central to NMDAR hypofunction in SCZ, dysfunctions in SR^-/-^ mice are likely spreading to other inhibitory cell-types including the SST and CCK neurons since the latter are expressing NMDARs (27, 28) as well. Additional studies are needed to elucidate the relative roles of the co-agonists at these different cell populations and the physiopathological relevance of these modulations in circuits dynamics of the deep cortical layers which control the top-down flow of information in the brain.

In summary, our study sheds new light on brain circuit’s physiology by uncovering the general principles of NMDARs regulation at a major class of GABAergic interneurons, the FS-PV^+^ INs, by upstream signals released in their surrounding microenvironment. Building on these findings, we identify that d-serine and glycine play complementary actions at somatodendritic NMDARs to control the excitatory drive and the GABAergic tonus with major outcomes for our understanding of the excitatory/inhibitory imbalance observed in SR^-/-^ mice. By doing so, we suggest that both are critical for network computations by engaging somatodendritic NMDARs of PV^+^-INs in a complex manner and that loss of their functions would recapitulate the synaptic deficits in disease like SCZ and could be exploited for the future development of more effective clinical interventions targeting inhibitory neurons.

## Materials and Methods

Detailed description of all methods and materials can be found in the SI Appendix.

All experiments were conducted in accordance with European and French directives on animal experimentation and with local ethical committee approval. Electrophysiological recordings were performed on 40 to 60 day-old (mean = 54 ± 10 days) male and female mice. All quantitative data are expressed as mean ± SEM. Statistical analyses performed are detailed in the SI Appendix and in figure legends.

## Supporting information

Main Manuscript

## Acknowledgments

We thank Zoltan Nusser (Institute of Experimental Medicine, Budapest), Dominique Debanne (UNIS, INSERM Aix-Marseille University) and Magalie Martineau for valuable discussion during the elaboration of the study, and for their critical reading and feedback on the manuscript. We also thank Prof Zoltan Nusser for having conducted high-freezing replica labelling of NMDARs on our demand and for sharing his results with us. Michael Rogawski and Sarah Denson (University of California, Davis) provided the SR knock-out mice. Laurent Venance (CIRB, College de France, Paris), Desdemona Fricker (INCC, University of Paris) and Maria-Cecilia Angulo (IPNP, Paris) provided the Ai9-tdTomato::PV-Cre transgenic line. The authors also acknowledge the assistance of Ayma Galland, Valérie Domergue and the personnel of the animal facility AnimEX in mouse breeding and care. We thank Maelle Rousselot for her technical assistance with immunostainings and microscopy sessions. This work was supported by the Fondation pour la Recherche Médicale (Equipe FRM DEQ20150331734 to J-PM), Centre National de la Recherche Scientifique (to J-PM), ENS Paris-Saclay (J-PM), Université Paris-Saclay (to J-PM), and by an EMBO Postdoctoral Fellowship (ALTF 981-2020 to INOS). LP was supported by a grant from Fondo di Ateneo per la Ricerca. Work in the laboratory of PS was supported by an SNSF Advanced Grant.

## Author Contributions

J-PM conceptualized and supervised the project. INOS, PL, SM, and BP carried out and analyzed the electrophysiological experiments. SM and J-PM performed confocal immunohistochemistry. ZO and PS supervised, conducted and analyzed FISH experiments. LP provided *Rg*DAAO and *Bs*GO. INOS and J-PM wrote the manuscript which was reviewed, edited and approved by all authors.

