## Supplementary material for "Principles of NMDA receptor co-agonism at cortical fast-spiking GABAergic interneurons in the adolescent prefrontal cortex": Main Manuscript

**This PDF file includes:**

Supporting text

Figures S1 to S13

SI References

Supporting Information Text

**SI Materials and Methods**

**Animals.** Experiments were performed on late adolescent (45 to 60 day-old; mean = 54 ± 10 days) male and female mice, with corresponding control littermates, from four different lines: C57BL/6; serine racemase knock-out (SR^-/-^) mice (C57BL/6 genetic background) (1) and parvalbumin(PV^+^)-*td*Tomato mice (Jackson Laboratories, stock no 027395), which express the red fluorescent protein *td*Tomato in PV-positive INs (C57BL/6 genetic background) (2). SR^-/-^ and SR^+/+^ mice were crossed with PV^+^-*td*Tomato animals to obtain double transgenic mice for ready identification of PV^+^-cells with either SR genotype. In some cases, to identify PV^+^ neurons, we used Ai9-tdTomato:PV-Cre transgenic mice generated from crossing PV-IRES-Cre knock-in mice (Jackson Laboratories, stock no. 008069) with Ai9-tdTomato reporter knock-mice (Jackson Laboratories, stock no. 007905). Only Tg/Tg::SR^+/+^ or Tg/Tg::SR^-/-^ were used in the study. Animals were housed in groups in polycarbonate cages and maintained on a 12/12 hr light/dark cycle (light on at 7am) in a temperature (22°C) and humidity-controlled room. Animals were given access to food and water *ad libitum*. All experiments performed in France complied with the European Union recommendations (2010/63/EU) and were approved by the French Ministry of Agriculture and Fisheries (authorization number authorization No. 75 1273) and by the University Paris-Saclay (accreditation number C92-019–01).

**Slice preparation**. Animals were deeply anesthetized with isoflurane before cervical dislocation and decapitation. The brain was then quickly removed and placed in ice-cold slicing artificial cerebrospinal fluid (ACSF) containing (in mM): KCl 3.5, MgSO_4_ 7, CaCl_2_ 0.5, D-glucose 11, sucrose 212, NaH_2_PO_4_ 1.2, NaHCO_3_ 26.2, L-ascorbic acid 1.2 and Na-pyruvate 2.4 (pH 7.3-7.4 and 302-313 mOsml/L), continuously bubbled with 95 % O_2_ / 5 % CO_2_. Acute coronal slices of the prefrontal cortex containing the prelimbic area (350 µm thickness) were then prepared for all mice lines using a vibrating blade microtome (Campden, UK) and allowed to recover in regular ACSF containing (in mM): NaCl 124, KCl 3.5, MgSO_4_ 1.5, CaCl_2_ 2.5, D-glucose 11, NaH_2_PO_4_ 1.2, NaHCO_3_ 26.2, saturated with 95 % O_2_ / 5 % CO_2_ (pH 7.4, 325 mOsml/L) during 30 min at 32 °C and for additional 30 min to 6 hours at room temperature before recordings.

**Brain slices electrophysiology acquisition and analysis.** Slices were transferred to a recording chamber mounted on an upright Slicescope Pro 6000 system (Scientifica, UK) and continuously perfused with oxygenated ACSF (2-3 mL/min) at near physiological temperature (30°-32°C, Heated perfusion tube HPT-2A, ALA Scientific Instruments Inc, USA). Recordings were performed from FS-PV^+^ INs located in layer 5 of the prelimbic region (PrL) of the medial PFC. Cells were visualized by infrared differential interference contrast and with a cooled digital CCD OrcaR^2^ Hamamatsu camera (C10600-10B, Hamamatsu Photonics, Japan) in combination with an on-line video contrast enhancement. Epifluorescence illumination (CoolLED pE-2 excitation system) was used to visualize *td*Tomato-positive interneurons (Ex/Em: 555/581nm). Recordings were obtained using a Multiclamp 700B amplifier (Molecular devices, USA), signals were filtered at 2 kHz and digitized at 5 kHz *via* a DigiData 1440A interface (Molecular Devices, USA). Data were collected and analyzed using pClamp10 software (Molecular Devices, USA). Series resistance and holding current were monitored throughout the experiment. Cells with an access resistance > 25 MΩ at resting potential were excluded from analyses as well as any cell for which a >20 % change in those parameters occurred during the course of the experiment.

**Intracellular solutions and control ACSF.** Electrophysiological recording were made using borosilicate glass pipettes (resistance = 3–5 MΩ) filled with either one of the two following intracellular solutions, depending on the experiment: K-gluconate solution (in mM: K-gluconate 120, KCl 10, HEPES 10, EGTA 0.2, MgATP 4.5, NaGTP 0.3 and Na-phosphocreatine 14, pH 7.2-7.3 ; 290-295 mOsm/L) or Cs-methanesulfonate solution (in mM: CsCH_4_O_3_S 140, CsCl 6, MgCl_2_ 2, HEPES 10, EGTA 1,1, QX-314 5 and MgATP 4, pH 7.2-7.3 ; 295-300 mOsm/L). In some experiments, MK801 (2-3 mM) was added to the intracellular solution to block NMDAR at the recorded cell. Neurons were first clamped at −70 mV and allowed to dialyze for 5 min before recording, or 10 min for experiments with intrapipette solution containing MK801. All recordings described in Figures 1-7 were done in the presence of strychnine (1 μM) and picrotoxin (50 µM) in the bathing ACSF to block glycinergic and GABA_A_ receptors, respectively. All drugs needed for control conditions were applied 10 minutes before recording and during the full length of the experiment. Subsequent drug effects were measured 5 minutes after a plateau was obtained and compared to baseline.

**Firing experiments.** Membrane and firing properties of PNs and FS-PV^+^ INs were recorded in current-clamp mode using K-gluconate solution including or not MK801. Results were obtained in response of square step current injections (500 msec) into the recorded cell from -100 pA to 200 pA with 20 pA steps unless specified. Spike frequency adaptation was measured as the ratio of the first inter-spike interval duration over the 8^th^ one at rheobase +40 pA.

**Evoked and spontaneous postsynaptic currents (PSCs).** Evoked PSCs were recorded in voltage-clamp mode and elicited by focal perisomatic extracellular stimulation using bipolar tungsten electrodes positioned at ~50 µm from the recording site and connected to a Digitimer DS3 Isolated Current Stimulus isolation unit (Digitimer, Ltd) at a frequency of 0.1 Hz unless specified otherwise. Evoked AMPA EPSCs were recorded from FS-PV^+^ INs clamped at -70 mV in the presence of d-AP5 (50 µM) in the bath while evoked NMDA EPSCs were isolated by maintaining FS-PV^+^ INs at -30mV or +40 mV in the presence of NBQX (20 μM) in the perfusion ACSF. For both cases, Cs-methanesulfonate intrapipette solution was used with or without MK801 (see Figure legends). The NMDAR/AMPAR ratio of the EPSCs was calculated as the ratio of peak amplitude of NMDA EPSC at +40 mV and AMPA EPSC at −70 mV. I–V relationship of NMDA EPSCs was measured at holding potentials from −70 mV to +40 mV with 10 mV steps and amplitude was normalized as a fraction of the amplitude at +40mV. NMDA input-output curves were obtained at +40 mV with extracellular stimulation between 10-50 μA. Evoked inhibitory PSCs (IPSCs) were recorded in voltage clamp mode and elicited by perisomatic extracellular stimulation as described for EPSCs at a frequency of 0.1 Hz unless specified otherwise. IPSCs were recorded on PNs clamped at -70mV using Cs-methanesulfonate intrapipette solution containing 2mM MK801. For spontaneous EPSCs recordings, from which tonic current measurements were taken, PNs were clamped at +40 mV in the presence of strychnine (1 μM), picrotoxin (50 µM) and NBQX (20 μM). For spontaneous IPSCs recordings, PNs were clamped at +10mV. All recordings of synaptic responses were made in the presence of CGP55845 (1 µM), DPCPX (100 nM), AM281 (250 nM) and (S)-MCPG (250 µM) in addition to strychnine and NBQX in bath ACSF to block GABA_B_Rs, A_1_Rs, CB_1_Rs and mGluRs and prevent network effects.

**Temporal summation and long-term potentiation.** FS-PV^+^ INs were recorded with the K-gluconate internal solution with QX-314 included when assessing long-term potentiation (LTP) to prevent firing of action potentials during induction (3). Temporal summation was investigated by two means: paired-pulse ratios (PPR) with inter-stimulus intervals (ISI) between 50-250 ms at a stimulus intensity of 50% the maximum response; and trains of 20 stimulations at varying frequencies (2-50 Hz). Excitatory LTP (eLTP) was investigated by maintaining FS-PV^+^ INs at -70mV during a 5 min baseline then at 0 mV during the conditioning stimulus. Afterward, the holding potential was set back at -70 mV for a 30 min post-conditioning recording. eLTP was elicited by a mild protocol consisting in high frequency tetanus (1 sec at 100 Hz repeated twice at a 20 sec interval) concurrently with postsynaptic depolarization of the FS-PV^+^ IN to 0 mV. PPR (ISI 30 ms) was recorded before baseline and 30 min after eLTP induction to inquire about the pre- versus post-synaptic locus of expression of the LTP. Inhibitory LTP (iLTP) was elicited by maintaining PN at -70mV and a theta burst stimulation (4 stimuli at 100Hz repeated 25 times at 5Hz), whose pattern mimics endogenous theta rhythms. As for eLTP, PPR was recorded before baseline and 30 min after iLTP induction (ISI 50 ms).

To further precise the locus of action of some drugs used in this study or the LTP locus, the coefficient of variation (CV) analysis method was used (4). The proportional change in the inverse square of the coefficient of variation (CV^–2^) was compared with the proportional change in the mean postsynaptic potential amplitude (M) to determine whether the quantal amplitude (q), the release probability (p) or the number of release sites (n) had changed. In a binomial distribution, CV^–2^ = [np/(1 – p)] and is therefore independent of q, while M = npq. When mean postsynaptic potential amplitude changes, no change in CV^–2^ indicates that only q has changed and a postsynaptic site is involved. A larger proportional change in CV^–2^ than in M indicates that p has changed, whereas a similar proportional change in the two parameters indicates that n has changed. Both presynaptic and postsynaptic sites are affected when a smaller proportional change in CV^–2^ than in M is revealed.

**Immunostainings and confocal microscopy.** Double transgenic mice (i.e PV^+^-*td*Tomato::WT and PV^+^-*td*Tomato::SR^-/-^) were anesthetized with intraperitoneal injection of sodium pentobarbital (60 mg/kg body weight) and perfused intracardiacally with phosphate buffered saline (PBS 0.1M ; pH7.4), followed by freshly dissolved 4% paraformaldehyde in PBS. The brains were then carefully removed and postfixed overnight in cold fixative. Brain coronal sections (25 µm thick) were obtained using a vibratome (Leica systems, VT1000S) collected in PBS and incubated for 30 min with PBS containing 0.5% sodium borohydride for quenching endogeneous fluorescence before blocking and permeabilisation with 0.3% Triton-X100 and 8% normal goat serum (NGS) in PBS for 1h. Brain sections were then stained 48h at 4°C with the single-domain antibody FluoTag-Q anti-RFP conjugated to AbberiorStar580 (Clone 1H7 ; NanoTag Biotechnologies) diluted at 1 :1000 in PBS containing 0.1% Tx-100 and 2% NGS. After five washes in PBS + 0.1% Tx-100, brain slices were in some case counterstained with DAPI (1 :10 000) for 5 min, before further washes and mounted in Vectashield (Vector Labs) to evaluate cortical lamination. Brain sections were examined with an inverted confocal laser-scanning microscope (Leica TCS SP8). Stacks of consecutive confocal images taken with a 10x or a 40x objective were acquired at 651 nm and 405 nm sequentially. Z projections were reconstructed using Fiji software.

**FISH experiments.** Brains were dissected from P14 or P50-55 male and female mice and snap-frozen in liquid nitrogen. Coronal sections (20 μm thick) were cut with a cryostat to include the prelimbic area (PrL) of the medial prefrontal cortex, adhered to Superfrost ultra plus slides (Thermo Scientific), and stored at −80 °C. Fluorescence *in situ* hybridization (FISH) was performed using the HCR RNA-FiSH (Molecular Instruments^TM^) protocol with the following modifications. Proteinase K was not applied to the sections, and the incubation with the hairpins was limited to 90 minutes to have single molecule resolution. The following probes were used in all hybridization reactions: Pvalb-B3 (commercially available, coupled to B3-488 hairpins), Grin1-B2 (commercially available, coupled to B2-546 hairpins). Grin2b, Grin2d, and Grin3a probes (B1) were custom-designed and coupled to B1-647 hairpins. DAPI was used to identify the nuclei. Images were acquired with an upright LSM700 confocal microscope (Zeiss) using a 40x Apochromat objective in z-stacks (19–22 images, 0.4 µm intervals), using laser lines of 405 nm (for exciting DAPI), of 488 nm (for Alexa 488), of 555 nm (for Alexa-546), and of 639 nm (for Alexa-647). PV-INs were identified based on the presence of the Pvalb probe signal, and the expression of NMDAR subunits was analyzed in PV-INs contained in layer 5 of PrL. A region of interest (ROI) was drawn to define the area of the cell soma. Thresholding for detection of the signal throughout the stacks and the number of pixels was done with Fiji software (https://fiji.sc/), using the TrackMate plugin (https://imagej.net/TrackMate). The number of puncta for each probe set was then counted with the TrackMate plugin in the ROI. The number of puncta in the ROI, normalized to the area, was used as a proxy for the expression strength of a given probe. To assess the effect of development on the ratio of the GluN subunits in PV^+^-INs, the number of puncta for Grin2b, 2d, and 3a were normalized to the number of Grin1puncta. Images were assembled using Fiji, OMERO (https://www.openmicroscopy.org/omero/), and Adobe Illustrator CS6 (https://www.adobe.com/uk/products/illustrator.html). Statistical tests for quantification of Grin1, Grin2b, Grin2d and Grin3a mRNA levels were performed in GraphPad Prism10 (https://www.graphpad.com/scientific-software/prism/). The number of dots per cell was averaged to yield an average expression level in P50-55 mouse for each Grin probe. This analysis was repeated in a total of N = 6 mice for each age group, and the statistical difference was tested with a non-parametric Mann-Whitney test for unpaired comparisons. Statistical difference for cumulative distribution of NMDAR subunits and the ratio of Grin2b, Grin2d to Grin1 mRNA levels was tested with non-parametric Kolmogorov-Smirnov test.

**Enzymes and drugs.** d-serine and glycine selective depletion was achieved by using recombinant wild-type *Rhodotorula gracilis* D-amino acid oxidase (*Rg*DAAO, EC 1.4.3.3) and recombinant H244K variant of *Bacillus subtilis* glycine oxidase (*Bs*GO, EC 1.4.3.19) respectively (5,6). *Rg*DAAO show a specific activity of 75 U/mg protein on d-serine, and variant *Bs*GO of 2.5 U/mg protein on glycine. Slices were first pre-incubated during 45 minutes at 32°C with either *Rg*DAAO or *Bs*GO (both at 0.4 U/mL) before recordings. Enzymes were then continuously bath-applied at 0.2 U/mL during the whole recording. 1-(2,4-Dichlorophenyl)-5-(4-iodophenyl)-4-methyl-N-4-morpholinyl-1H-pyrazole-3-carboxamide (AM281), N-[3-(4′-Fluorophenyl)-3-(4′-phenylphenoxy)propyl]sarcosine hydrochloride (ALX5407), (2S)-3-[[(1S)-1-(3,4-Dichlorophenyl)ethyl]amino-2-hydroxypropyl](phenylmethyl)phosphinic acid hydrochloride (CGP55845), d-(2R)-amino-5-phosphonovaleric acid (d-AP5), γ-d-glutamylglycine (γ-DGG), 8-Cyclopentyl-1,3-dipropylxanthine (DPCPX), QX314 bromide, 4-((1H-Indol-7-yl)carbamoyl)phenyl diethylcarbamate (NAB-14), NBQX disodium salt, MK801 maleate, N-methyl-D-aspartate (NMDA), picrotoxin, Ro 25-6981 and (S)-α-Methyl-4-carboxyphenylglycine ((S)-MCPG) were from either Tocris (Bristol, UK), or HelloBio (Bristol, UK). 5-Chloro-benzo[d]isoxazol-3-ol (CBIO) was from Maybridge (Cornwall, UK). d-serine, strychnine and 5,7-dichlorokynurenic acid (DCKA) were from Sigma-Aldrich France (Saint-Quentin, France). ω-AgaTx IVB, ω-conotoxin and tetrodotoxin were from Alomone labs Ltd (Jerusalem, Israel), dissolved in water and stored at -20 °C. When drugs were dissolved in DMSO, final concentration of the vehicle in bathing ACSF was kept at 1/4000.

**Analysis and statistics.** Analysis was conducted using Prism 8 software (GraphPad, San Diego, California). Results are expressed as mean ± SEM and a p-value lesser than 0.05 was considered significant. In the figures legends, n refers to the number of recorded cells sampled from 2-6 mice depending on conditions. Two-way ANOVA with repeated measurements were done for firing frequency, input/output curve, NMDA IV curve and short-term plasticity experiments with post-hoc multiple comparison tests using a Bonferroni-Dunn correction factor when necessary. For time-course experiment and LTP, the mean amplitude of the last 5 minutes of recording (expressed in percent of change compared to control) was tested versus 0 (and versus 100 for LTP) with one-sample Wilcoxon test. Other comparisons were done using the Wilcoxon test for paired data and the Mann-Whitney test for unpaired data. To further precise the locus of action of the drugs, the coefficient of variation (CV) analysis method was used (4).

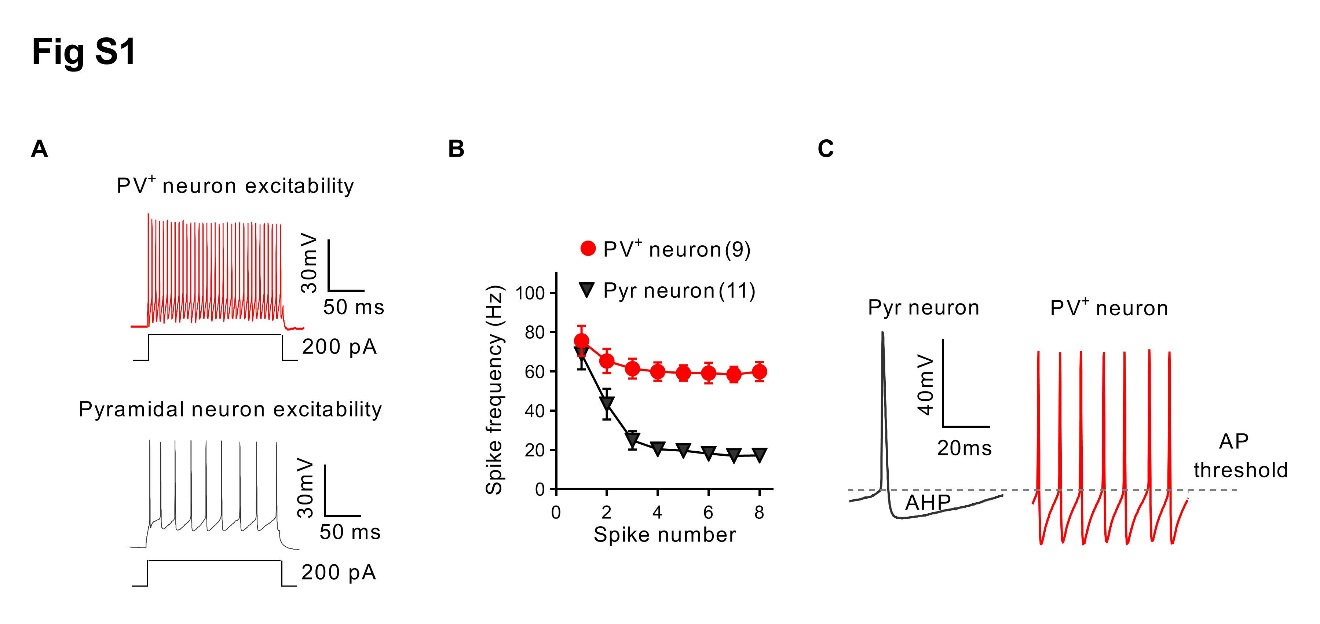

Fig. S1. Firing profiles of pyramidal cells and FS-PV^+^ interneurons from the layer 5 of mPFC. (A) Representative traces of the firing pattern for a FS-PV^+^ interneurons (red) and regular spiking pyramidal cells (Pyr, black) in response to a 200pA depolarizing current step. (B) Frequency adaptation is limited in FS-PV^+^ interneurons while present in Pyr cells. (C) Comparison of spikes in Pyr neuron (black trace) and FS-PV^+^ interneuron (red trace) highlighting the faster kinetic and bigger after-hyperpolarisation characteristic of spikes from FS-PV^+^ neurons.

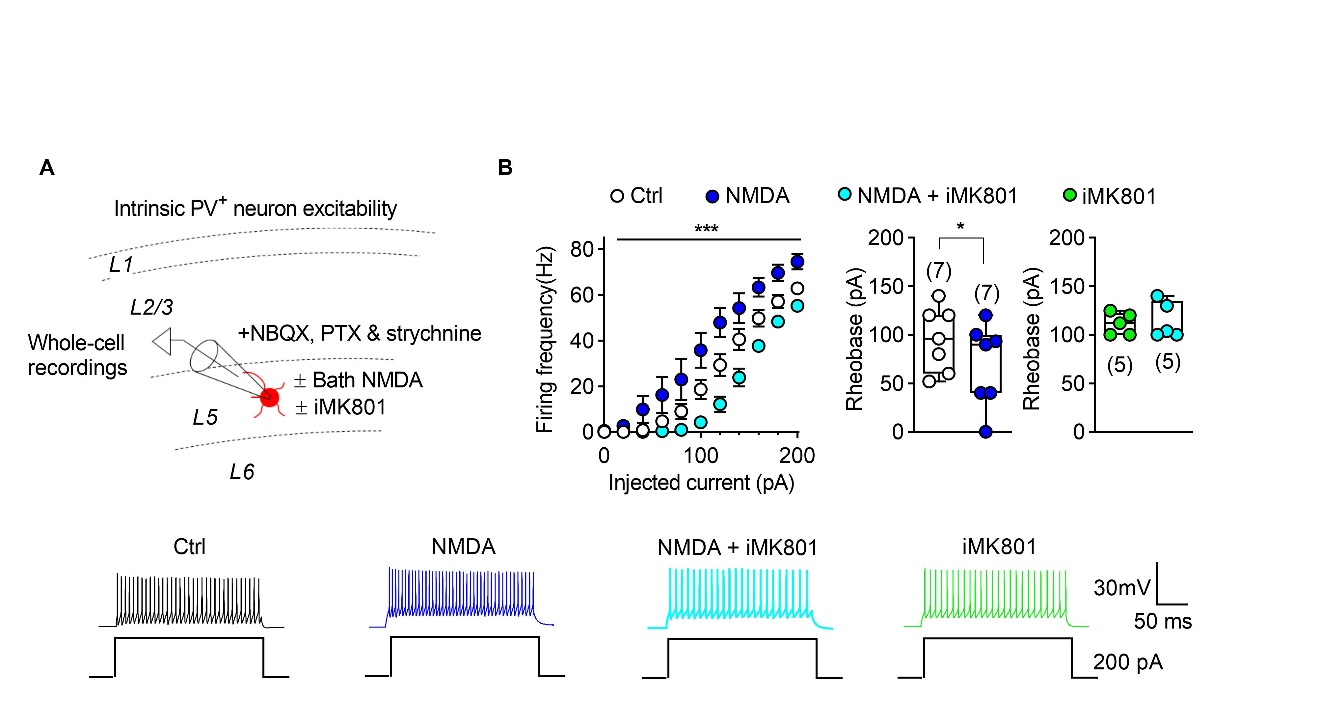

**Fig. S2. NMDAR are present on adolescent FS-PV^+^ interneurons and control their excitability**. (**A**), Experimental protocol for recording intrinsic excitability of FS-PV^+^ interneurons from PV^+^-tdTomato mice. (**B**) *Top left,* Activation of NMDAR by application of NMDA (dark blue) enhanced firing frequency while their blockade by intra-pipette MK801 (iMK801, cyan) had the opposite effect (Two-way ANOVA, ****p=0.0004*, F_(2,27)_=10,72).*Top right,* NMDA application decreased rheobase (Wilcoxon test, ***p=0.0313*) while iMK801 increased it (Mann-Whitney test, **p=0.0180*). iMK801 suppressed the effect of NMDA on firing frequency (left, Two-way ANOVA, *p=0.8093*) and on rheobase (right, Wilcoxon test, *p=0,7500*). *Bottom,* representative traces of the firing pattern of FS-PV^+^ interneurons for all tested conditions.

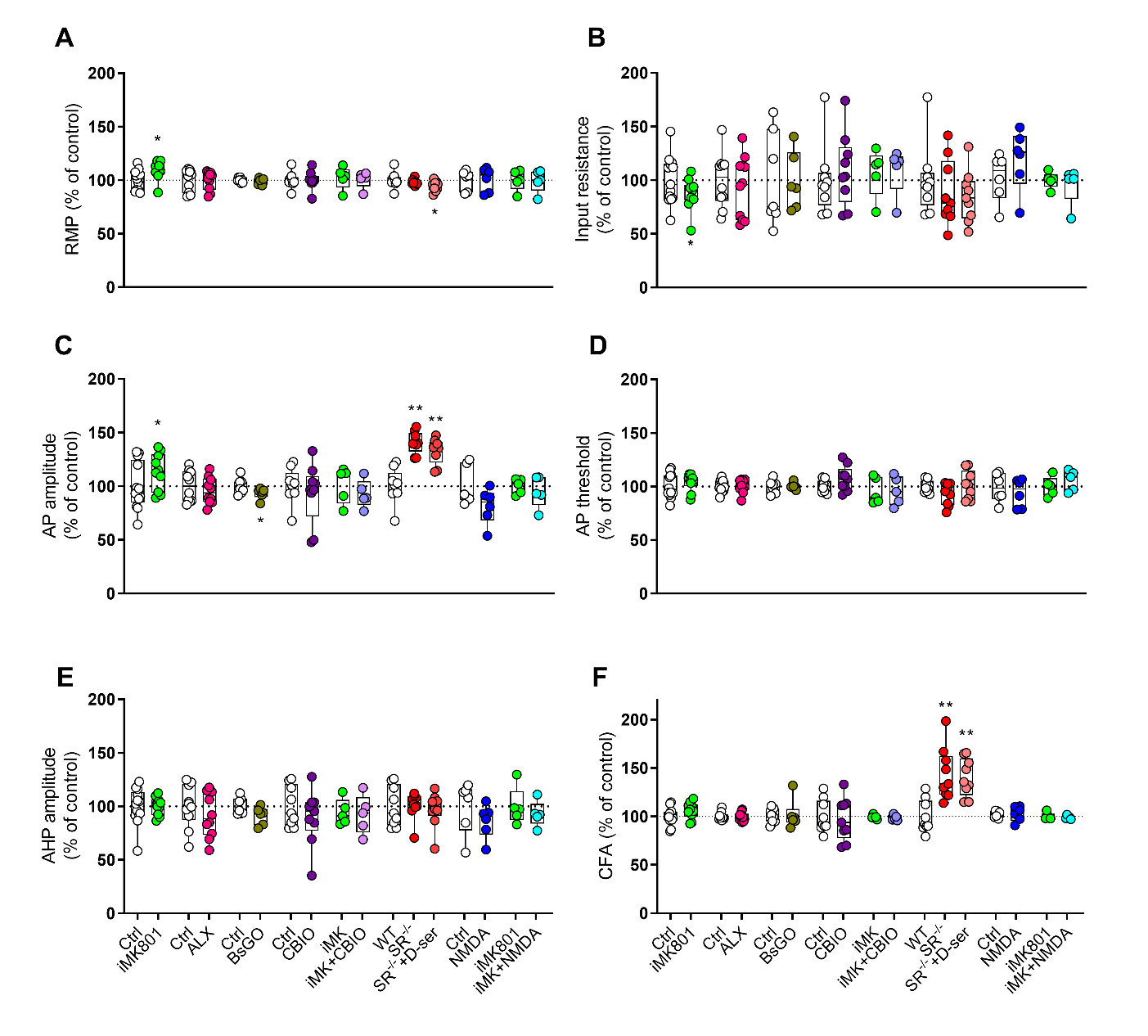

**Fig. S3. Effects of gain- and loss-of-function of d-serine on membrane properties of FS-PV^+^ interneurons.** Boxplot representing percentage of control values for resting membrane potential (RMP) (**A**), input resistance (**B**), action potential amplitude (AP amplitude) (**C**), action potential threshold (AP threshold) (**D**), after-hyperpolarization amplitude (AHP amplitude) (**E**) and coefficient of frequency adaptation (CFA) (**F**), respectively, on different conditions depicted in Figures 1, 2, 4 and S2. Initial iMK801 group showed significant increases in RMP (**p=0.0273*) and AP amplitude (**p=0.0371*) with a significant decrease in input resistance (**p=0.0273*), but such changes were not seem in other iMK801 groups. SR^-/-^ groups showed significant increases in AP amplitude (***p=0.0039* to both) and CFA (***p=0.0039* to both). Wilcoxon test *vs* 100.

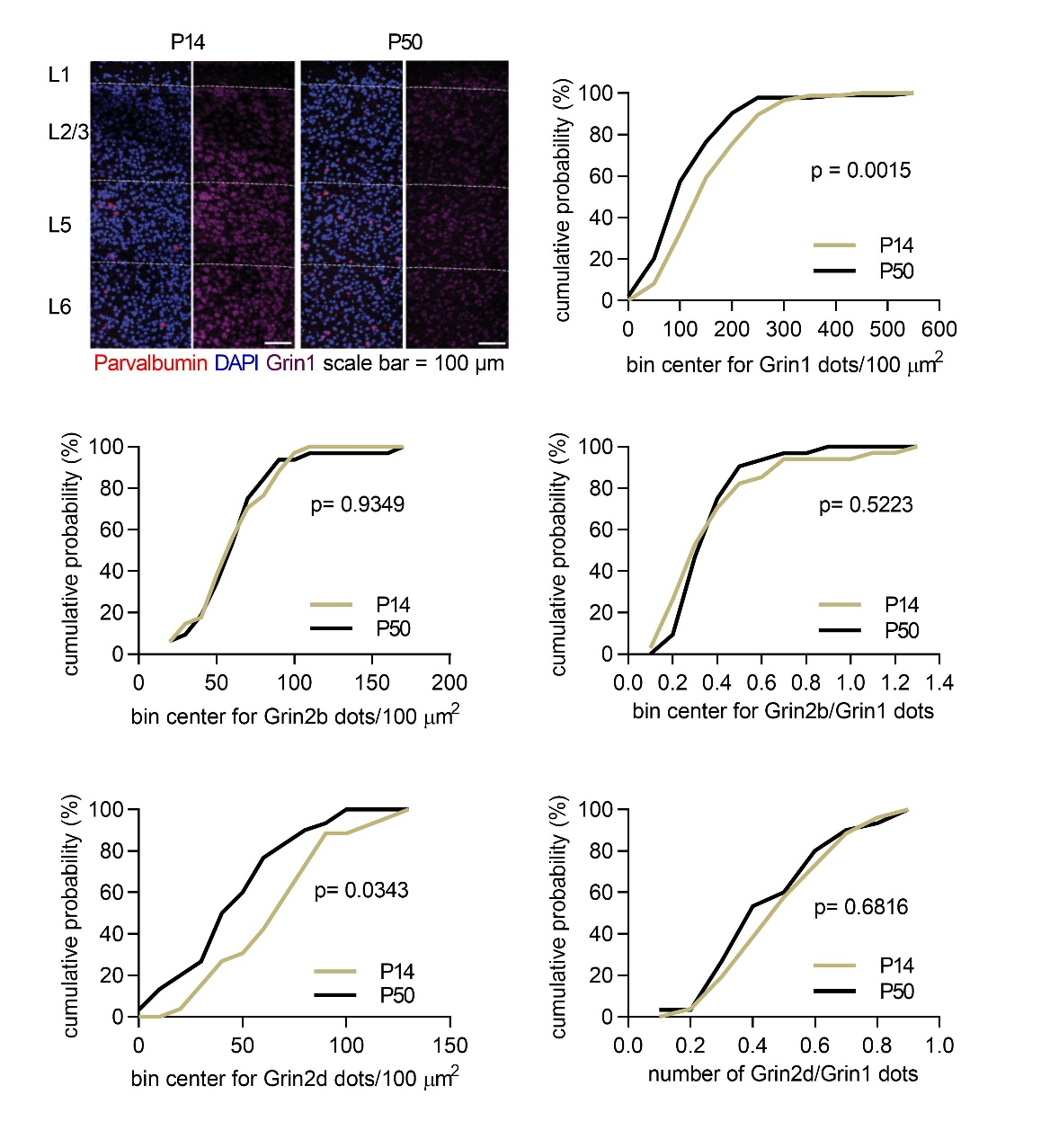

**Fig. S4. Developmental regulation of NMDAR subunit mRNA expression in parvalbumin interneurons of the medial prefrontal cortex.** Representative fluorescence in situ hybridization (FISH) images showing expression of Pvalb (red), Grin1 (magenta), and DAPI (blue) in the prelimbic cortex of juvenile (P14) and late-adolescent (P50) mice. PV interneurons were identified by Pvalb expression, and mRNA puncta were quantified within the soma. Scale bar, 100 μm. Right panels show cumulative distributions of Grin1, Grin2b, and Grin2d mRNA puncta density (dots/100 μm²) as well as the normalized Grin2b/Grin1 and Grin2d/Grin1 ratios in PV interneurons at P14 and P50. Grin1 expression decreased significantly with development (Kolmogorov–Smirnov test, *p* = 0.0015), whereas Grin2b expression and the Grin2b/Grin1 ratio were unchanged (*p* = 0.9349 and *p* = 0.5223, respectively). Similar to Grin1, Grin2d expression decreased significantly during development (*p* = 0.0343), while the Grin2d/Grin1 ratio showed a similar trend that did not reach statistical significance (*p* = 0.0816). Quantification was performed from layer 5 PV interneurons in the prelimbic cortex of P14 and P50–55 mice as described in the methods.

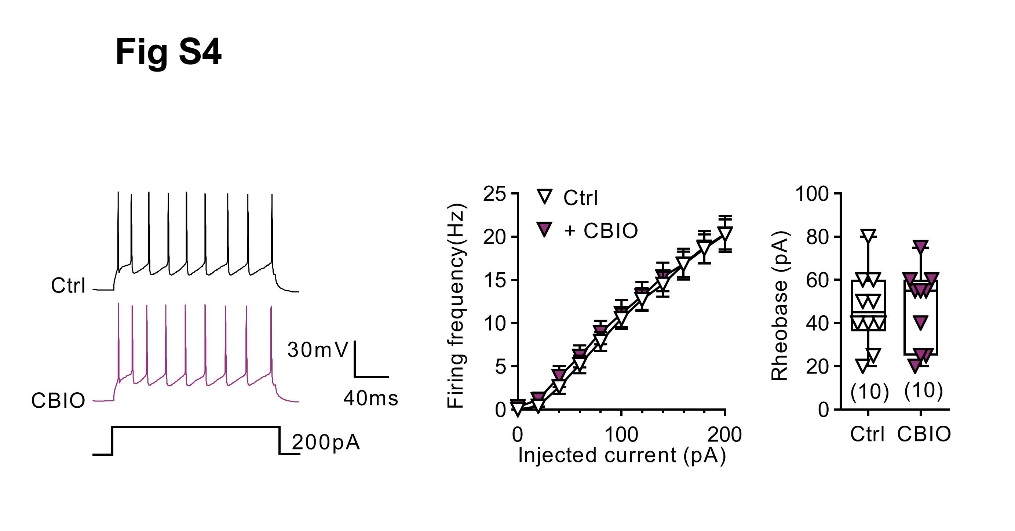

**Fig S5. Firing profile of pyramidal cells is not affected when d-serine levels are potentiated with CBIO.** *On the left*: representative traces of the firing pattern of Pyr cells under control conditions (black trace) and after application of the DAAO inhibitor CBIO (1µM) (purple traces). Elevation of d-serine extracellular levels by CBIO affected neither firing frequency (*in the middle*, two-way ANOVA, p=0.2565, n=11) nor rheobase (*on the right*, Wilcoxon test, p>0.9999, n=11).

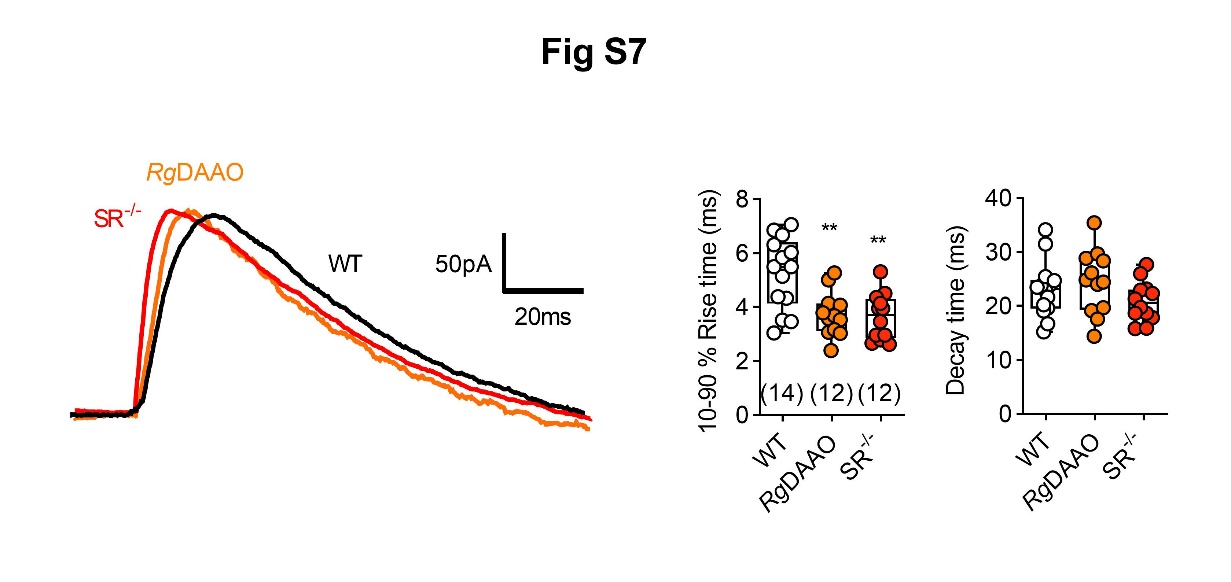

**Fig. S6. Kinetic of NMDA-EPSCs is altered when d-serine is absent**. NMDA-EPSCs were recorded at +40 mV. Left, averaged traces in wild-type before (black trace) and after treatment of *Rg*DAAO (orange) and in SR^-/-^ (red). Right, boxplots show analysis of activation and deactivation constants. The rise time constant of NMDA-EPSCs was reduced when WT slices were treated with *Rg*DAAO (*Rg*DAAO, n= 12 cells *vs* WT, n=14 cells, *p*=0.0057, Mann-Whitney test) or from SR^-/-^ mice (SR^-/-^, n= 12 cells *vs* WT, n=14 cells, *p*=0.0017, Mann-Whitney test) while decay time constant remained unaffected. ***p*< 0.01. Data are expressed as mean ± SEM

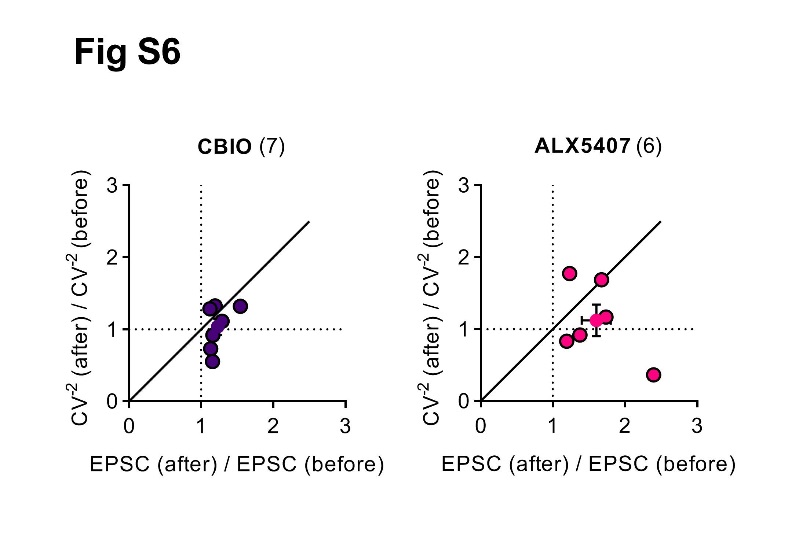

**Fig. S7. Postsynaptic modulation of NMDA-EPSCs by d-serine and glycine.** Plotting the ratio (after/before CBIO or ALX5407 bath application) of the inverse square of the coefficient of variation (CV^-2^) against the ratio (after/before CBIO or ALX5407 bath application) of the amplitude shows that for both compounds the CV^-2^ did not change (One sample Wilcoxon test *vs* 1: *p=0.6875* for both CBIO (1.03 ± 0.11, n=7) and ALX5407 (1.12 ± 0.22, n=6). On the contrary, the amplitude ratio was significantly higher than one (One sample Wilcoxon test *vs* 1: *p=0.0156* for CBIO (1.23 ± 0.06, n=7) and *p=0.0313* for ALX5407 (1.60 ± 0.18, n=6).

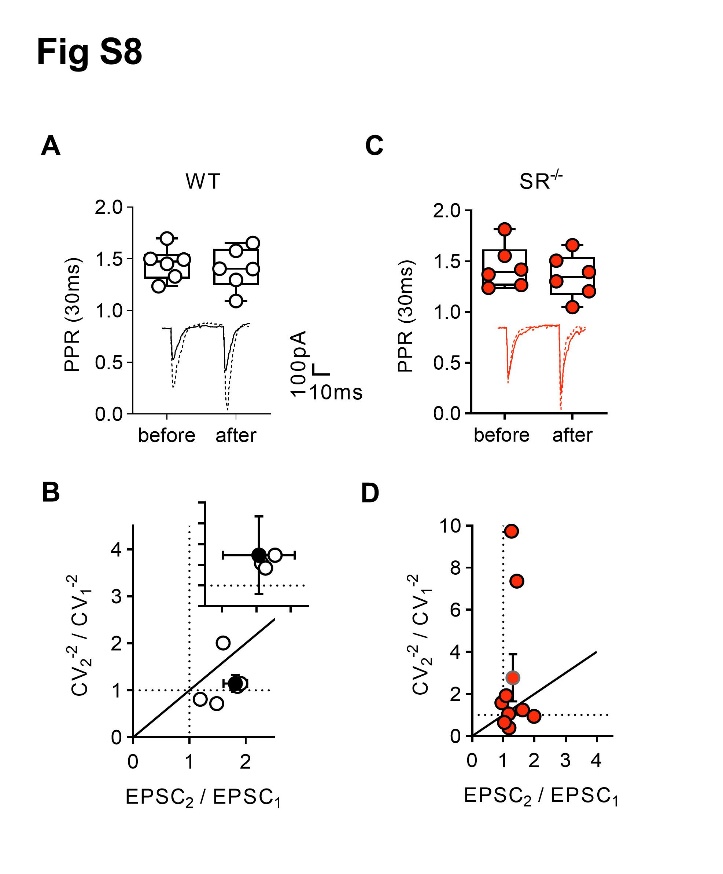

**Fig. S8. PPR and CV analysis of the tetanus-induced eLTP at the PN-*to*-PV^+^ synapse. (A and C)** Paired-pulse ratio in control (1.451 ± 0.065 *vs* 1.405 ± 0.082, Wilcoxon test *p=0.8438*, n=6) and SR^-/-^ (1.444 ± 0.088 *vs* 1.353 ± 0.089, Wilcoxon test *p=0.6875*, n=6) groups were similar before and after LTP. Insets show representative EPSCs separated by 30 ms for control (before eLTP, continuous black trace and after eLTP black dashed trace) and SR^-/-^ groups (before eLTP, continuous red trace and after eLTP, red dashed trace). (**B and D)** Plotting the ratio (after/before LTP induction) of the inverse square of the coefficient of variation (CV^-2^) against the ratio (after/before LTP induction) of the amplitude revealed a non-significant increase of CV^-2^ for both groups (One sample Wilcoxon test *vs* 1: *p=0.0625* for Ctrl (2.47 ± 0.58, n=6) and *p=0.2031* for SR^-/-^ (2.77 ± 1.12, n=9). On the contrary, the amplitude ratio was significantly higher than one (One sample Wilcoxon test *vs* 1: *p=0.0313* for Ctrl (1.83 ± 0.22, n=6) and *p=0.0117* for SR^-/-^ (1.31 ± 0.11, n=9).

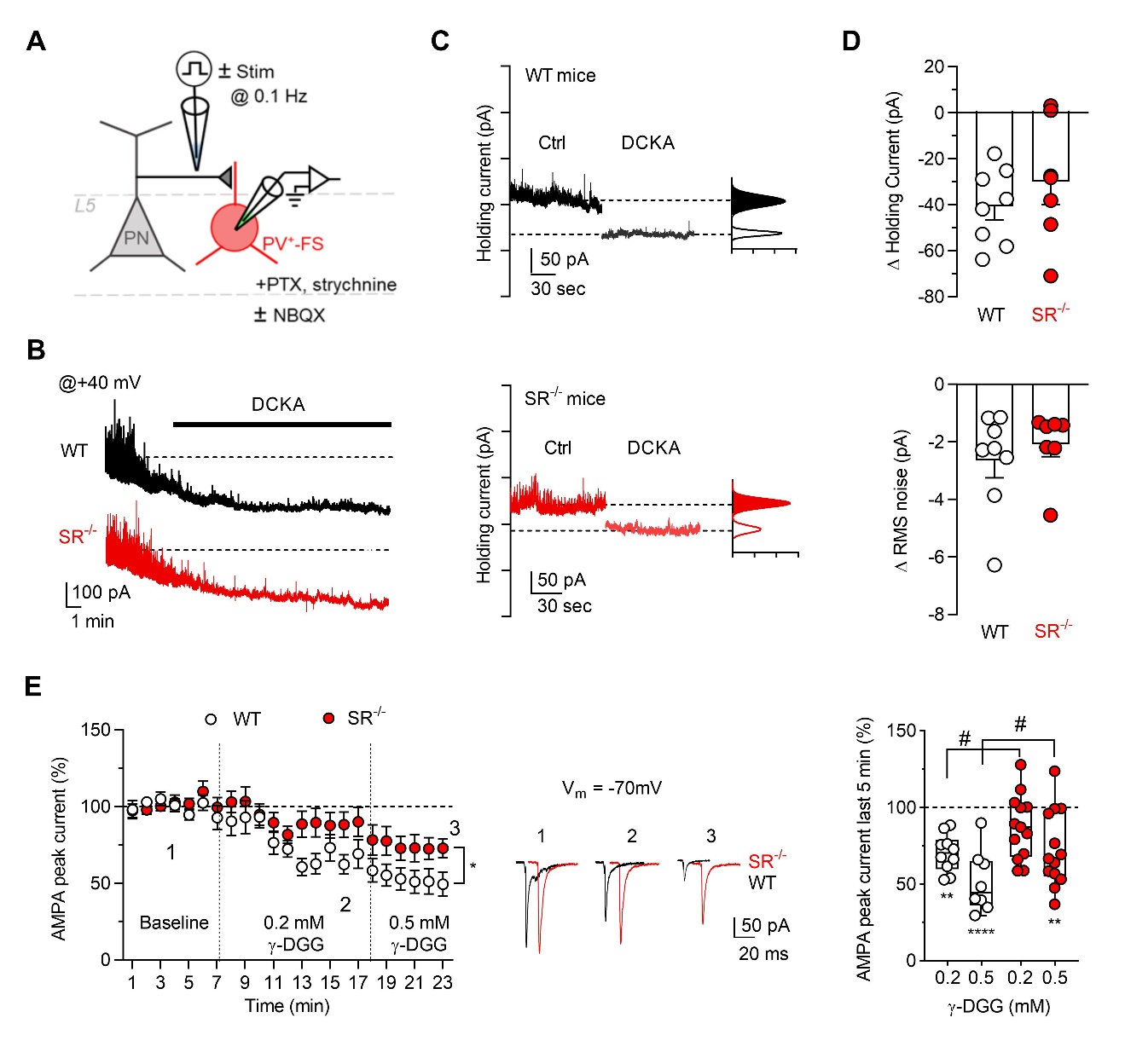

**Fig S9. Chronic d-serine depletion modifies tonic NMDAR control of FS-PV^+^ interneurons.** **(A)** Schematic diagram of recording electrode position in mPFC layer 5, where FS-PV^+^ INs were recorded in PV^+^-*td*Tomato::WT or PV^+^-*td*Tomato::SR^-/-^ mice. Recordings were made with bath application of PTX, strychnine (all panels) and NBQX (panels B to D) for isolation of NMDAR activity. Tonic current and sEPSCs recordings (panels B to E) were done at +40 mV while γ-DGG recordings (panel F) were done at -70mV. **(B to D)** Shift in the holding currents induced by DCKA (50 µM) in WT (white, n= 8 cells) and SR^-/-^ (Red, n=7 cells) mice. DCKA was bath applied during 20 min. Example traces, time course, and summary plot of the variations in the holding currents in WT and in SR^-/-^ FS-PV^+^ INs upon DCKA. Holding current and noise reduction induced by DCKA: *p*= 0.4634, Mann-Whitney test. **(E)** Effect of γ-DGG, a fast-dissociating AMPA antagonist on AMPA EPSCs at two concentrations (0.2 and 0.5 mM) recorded in WT (white symbols and black traces) and in SR^-/-^ (red). Time course (left) shows the inhibitory action of γ-DGG (*p*= 0.0209, two-way ANOVA with repeated measurements). Examples traces are shown. Box plots (right) representing the mean average changes induced by γ-DGG at the two doses in WT and in SR^-/-^ mice. Comparisons with Ctrl for each dose across genotype were done using one sample Wilcoxon test. 0.2 mM WT (n=10 cells) and 0.5 mM WT (n=8 cells) *vs* Ctrl WT (n= 10 cells), *p*=0.002 and *p*=0.0078, respectively. 0.2 mM SR^-/-^ (n=13 cells) and 0.5 mM SR^-/-^ (n=13 cells) *vs* Ctrl SR^-/-^ (n= 13 cells), *p*=0.0681 and *p*=0.0024, respectively. Comparisons across genotype for each dose were done using Mann-Withney test. 0.2 mM WT *vs* 0.2 mM SR^-/-^, *p*= 0.0493; 0.5 mM WT *vs* 0.5 mM SR^-/-^, *p*= 0.0446. *,^#^*p*< 0.05, ***p*< 0.01, ****p*< 0.001, *****p*< 0.0001. Data are expressed as mean ± SEM.

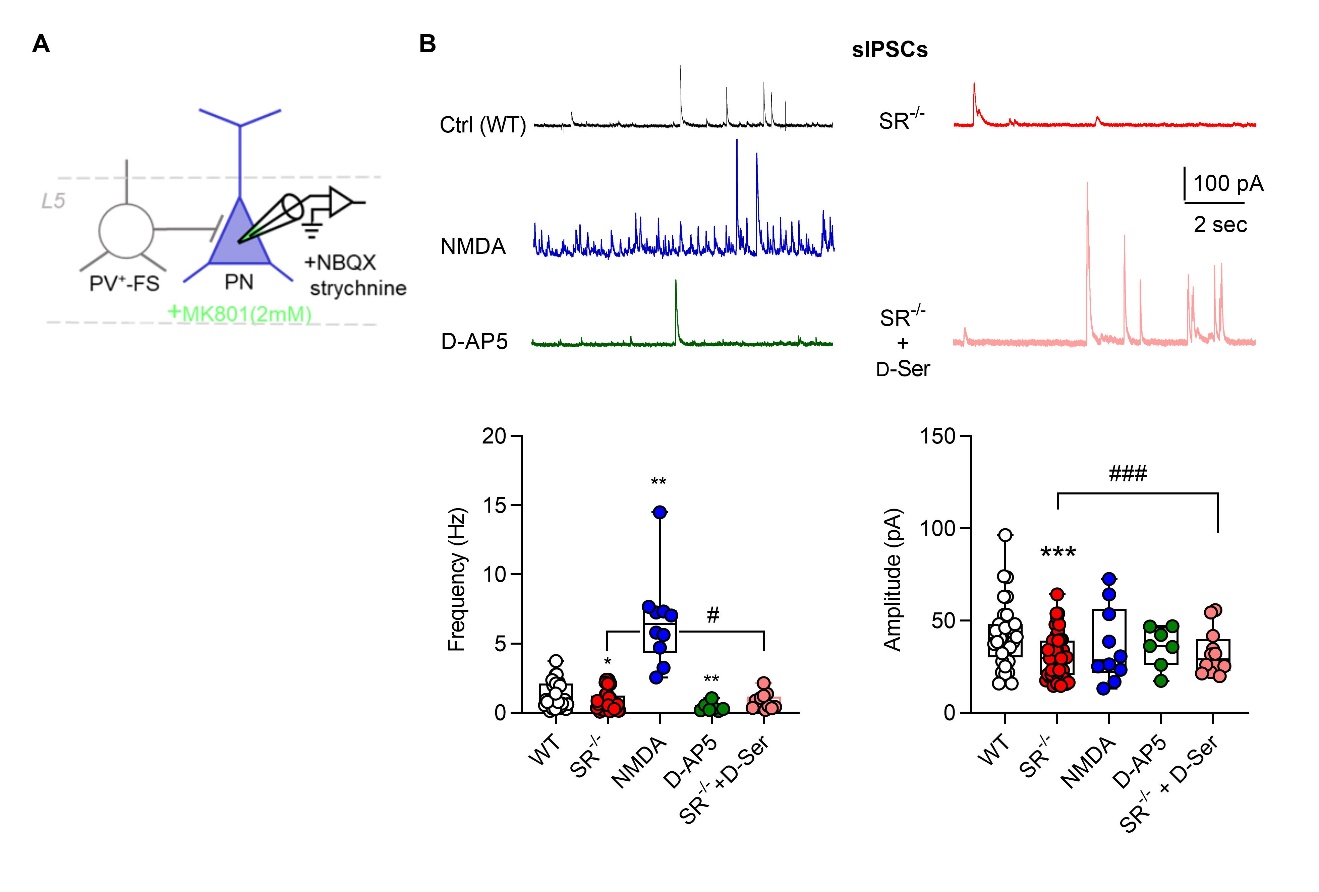

**Fig S10. Effect of NMDARs modulation on spontaneous inhibitory postsynaptic currents.** Recordings were made for at least 30 min with MK801 included in the patch pipette for blocking the NMDAR located at the PN and with the PN neuron clamped at +10 mV. **(A)** Schematic diagram showing the position of the recording electrode in mPFC layer 5, where spontaneous synaptic GABA_A_ currents at pyramidal neurons (PN) were recorded in WT or SR^-/-^ mice. **(B)** Upper, 10 sec samples traces are represented for each condition. Box plots (below) show the averaged changes for the frequencies and amplitudes of sIPSCs. NMDA (dark blue) increased sIPSCs frequency but not amplitudes (n= 33 Ctrl and 10 NMDA cells, *p*< 0.0001, Mann-Whitney test) while D-AP5 (green) decreased frequencies (n= 33 Ctrl and 7 D-AP5 cells, *p*=0.0074, Mann-Whitney test). sIPSCs frequencies and amplitudes in SR^-/-^ mice (red, n=41 cells) were reduced (*p=0.0438* for frequencies, *p*= 0.001 for amplitudes, Mann-Whitney test) when compared to WT mice. Bath-applied 20 µM d-serine (pink, n=12 cells) reversed the deficits observed on the frequencies (*p=0.0093* *vs* before d-Ser, and amplitude (*p=0.001 vs before d-Ser*) in SR^-/-^ mice. *^,#^*p*<0.05, ***p*<0.01, ^***,###^*p*< 0.001. Data are expressed as mean ± SEM

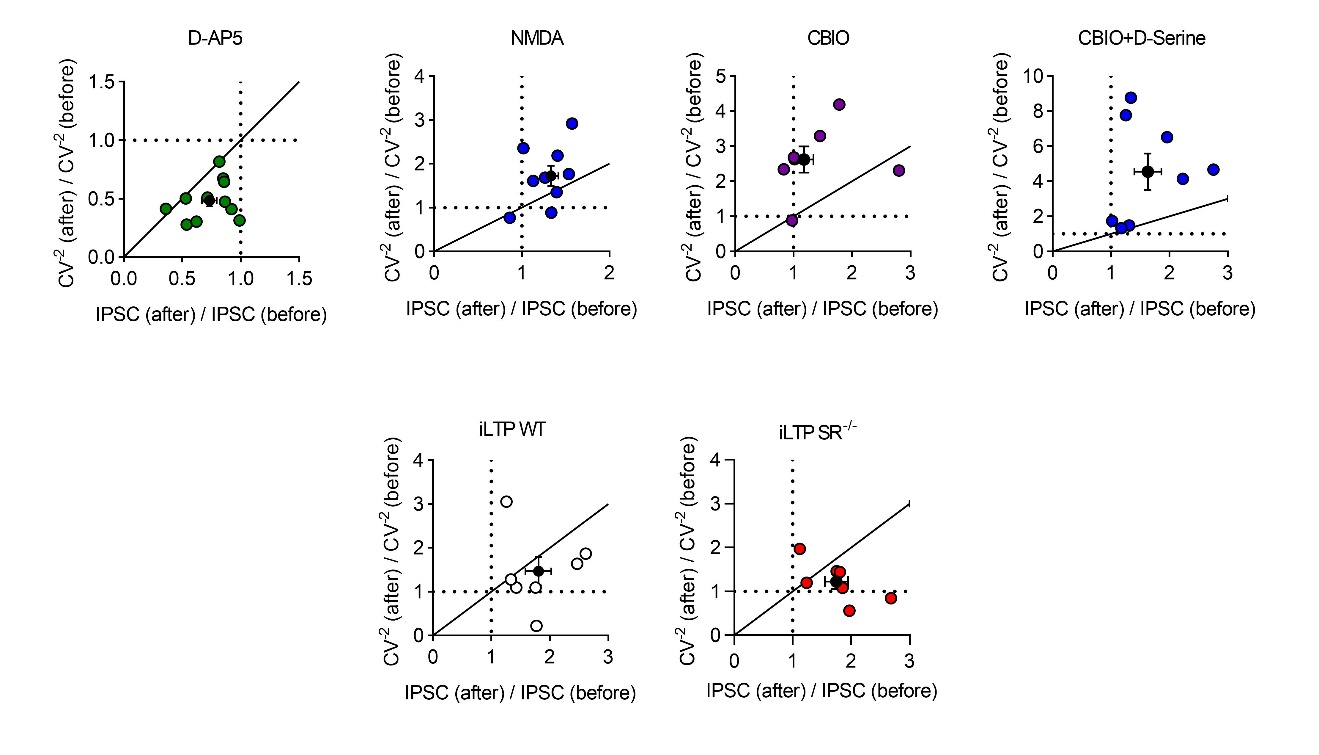
**A**

**B**

**Fig. S11. Variance analysis posits the presynaptic locus of the effect of drugs on IPSCs and the postsynaptic locus of iLTP. A)** Plotting the ratio (after/before D-AP5, NMDA, CBIO or CBIO + d-serine bath application) of the inverse square of the coefficient of variation (CV^-2^) against the ratio (after/before drug application) of the amplitude shows that for each condition the CV^-2^ was significantly different than one (One sample Wilcoxon test *vs* 1), indicating a presynaptic effect in each group. The CV^-2^ ratio was significantly lower than 1 for the D-AP5 treated group (0.49 ± 0.05, p=0.0010, n=11) and significantly higher than 1 for the NMDA (1.72 ± 0.23, p=0.0195, n=9), CBIO (2.62 ± 0.38, p=0.0313, n=7 ) and CBIO+D-Serine (4.55 ± 1.03, p=0.0078, n=8). Likewise, the amplitude ratio was significantly (One sample Wilcoxon test *vs* 1) lower than 1 for the D-AP5 treated group (0.73 ± 0.06, p=0.0010) and higher than 1 for the NMDA (1.28 ± 0.08, p=0.0195) and the CBIO+D-serine (1.63 ± 0.22, p=0.0078) treated groups. Of note, the CBIO treatment did not increase significantly the amplitude ratio (1.41 ± 0.26, p=0.2969). **B)** Plotting the ratio (after/before iLTP induction) of the inverse square of the coefficient of variation (CV^-2^) against the ratio (after/before iLTP induction) of the amplitude revealed a non-significant change in CV^-2^ for both groups (One sample Wilcoxon test *vs* 1: p=0.1563 for WT (1.46 ± 0.33, n=7) and p=0.2969 for SR^-/-^(1.22 ± 0.17, n=7). On the contrary, the amplitude ratio was significantly higher than 1 for both groups (One sample Wilcoxon test *vs* 1: p=0.0156 for WT (1.81 ± 0.20, n=7) and p=0.0156 for SR^-/-^(1.78 ± 0.19, n=7).

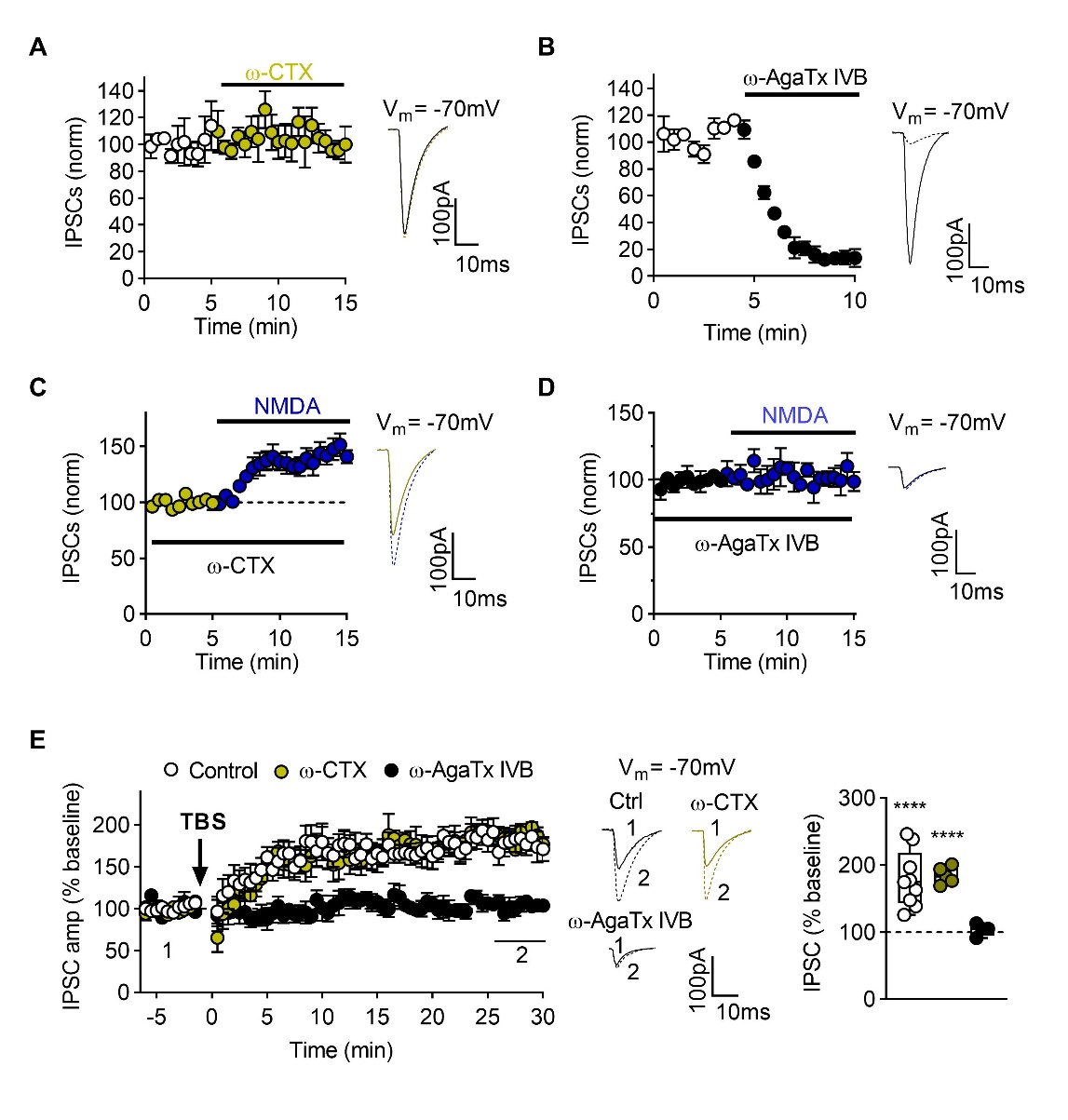

**Fig. S12. CCK^+^ interneurons are not involved in IPSCs elicited by perisomatic electrical stimulations.** All recordings were made at -70 mV with MK801 infused in the patch pipette to block NMDARs at PNs. **(A-B)** ω-conotoxin, which inhibits GABA release from CCK^+^ does not affect the IPSCs (n=5 cells) while ω-agatoxin IVB which blocks release of GABA by PV^+^-INs nulls IPSCs recorded at PNs (n=4 cells). **(C-D)** ω-conotoxin does not prevent the potentiating effect of bath-applied NMDA (10 µM) while ω-agatoxin IVB does. **(E)** Theta Burst stimulation was used to induced LTP at the inhibitory PV^+^- *to* PN synapse. Left, Time-course shows the iLTP in control, under ω-conotoxin or ω-agatoxin IVB. Right, mean normalized amplitude of the 5 last minutes for each group indicated that iLTP is normally induced in control (n= 9 cells, *p*< 0.0001, Wilcoxon test *vs* 100) and under ω-conotoxin (n= 5 cells, *p*<0.0001, Wilcoxon test *vs* 100) but is absent when ω-agatoxin IVB is applied (n= 4 cells, *p*= 0.3212, Wilcoxon test *vs* 100) condition. *****p*< 0.0001. Data are expressed as mean ± SEM

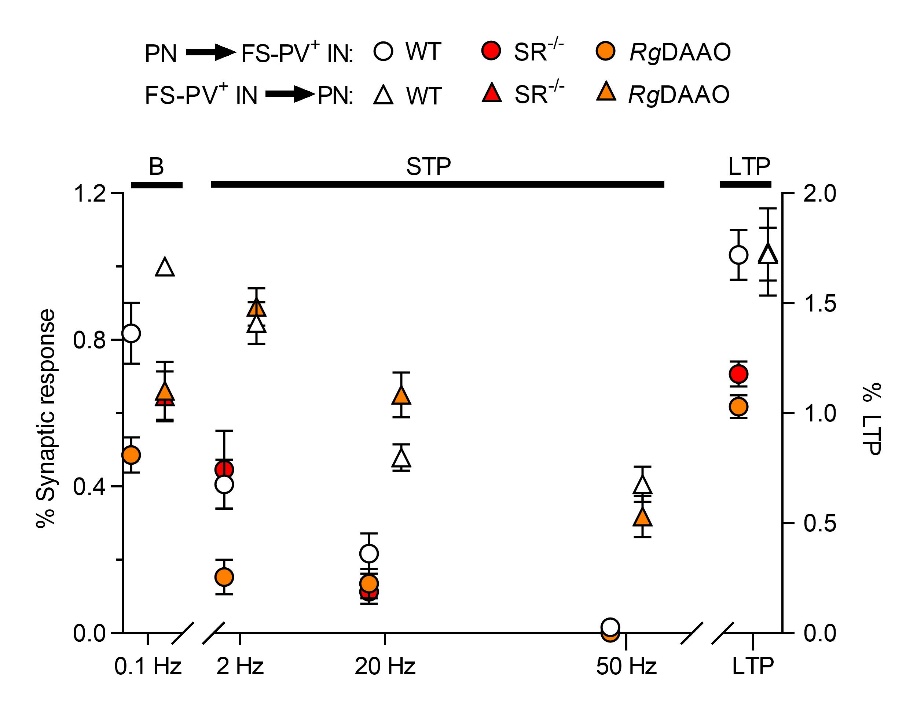

**Fig S13. Summary plot of the changes induced on EPSCs and IPSCs by varying the synaptic activity regime.**
